# Mechanical cell competition in heterogeneous epithelial tissues

**DOI:** 10.1101/869495

**Authors:** R. J. Murphy, P. R. Buenzli, R. E. Baker, M. J. Simpson

## Abstract

Mechanical cell competition is important during tissue development, cancer invasion, and tissue ageing. Heterogeneity plays a key role in practical applications since cancer cells can have different cell stiffness and different proliferation rates than normal cells. To study this phenomenon, we propose a one-dimensional mechanical model of heterogeneous epithelial tissue dynamics that includes cell-length-dependent proliferation and death mechanisms. Proliferation and death are incorporated into the discrete model stochastically and arise as source/sink terms in the corresponding continuum model that we derive. Using the new discrete model and continuum description, we explore several applications including the evolution of homogeneous tissues experiencing proliferation and death, and competition in a heterogeneous setting with a cancerous tissue competing for space with an adjacent normal tissue. This framework allows us to postulate new mechanisms that explain the ability of cancer cells to outcompete healthy cells through mechanical differences rather than by having some intrinsic proliferative advantage. We advise when the continuum model is beneficial and demonstrate why naively adding source/sink terms to a continuum model without considering the underlying discrete model may lead to incorrect results.

## 1 Introduction

In cell biology, epithelial tissues are continuously experiencing forces and replacing cells, through cell proliferation and death, to maintain homeostasis. These tissues can be naturally heterogeneous or heterogeneous due to to cancer development and progression (Han et al. 2019, Plodinec et al. 2012). This heterogeneity is observed at multiple scales, from sub-cellular to cellular to the tissue scale (Trepat et al. 2018), and can result in cell competition. Cell competition can act as a quality control mechanism in tissue development or as a defence against precancerous cells, and harnessing cell competition has been suggested as a possible approach to enhance both cell-based cancer and regenerative therapies (Powell 2019). Therefore, gaining a greater understanding of the mechanisms underlying cell competition is very desirable. In mathematical models of cell competition the classical hypothesis is that cells compete due to differences in their intrinsic proliferation rates. However, this may not be true and mathematical models have began to explore the role of different mechanisms (Lee et al. 2017). We will explore mechanical cell competition.

In the emerging field of mechanical cell competition, *winner* cells compress neighbouring cells promoting tissue crowding and regions of higher density, which leads to cell death (Bras-Pereira et al. 2018, Levayer 2019, Wagstaff et al. 2016), while cell proliferation occurs in regions of lower density (Gudipaty et al. 2017). In this work, we focus on mechanical cell competition arising from the coupling of a cell-based model of epithelial tissue mechanics with cell-length-dependent proliferation and death mechanisms. We consider mechanical forces to be driven by cell stiffness which is important for cancer progression (Samuel et al. 2011), cancer detection (Plodinec et al. 2012), morphogenesis (Fletcher et al. 2017), and wound healing (Evans et al. 2013). A grand challenge in cell biology is to understand how tissue-level outcomes are related to cell-based mechanisms, especially when experiments are performed by focusing on a single scale, and many cellular processes occur over multiple overlapping timescales (Cadart et al. 2019, Wyatt et al. 2016). Therefore, we apply mathematical modelling with *in silico* simulations to develop a framework to quantitatively connect cell-level mechanisms with tissue-level outcomes.

Many mathematical modelling frameworks, including both discrete models and continuum models, have been used to study cell migration and cell proliferation. In discrete models individual cell properties and inter-cellular interactions can be prescribed (Osborne et al. 2017, Pathmanathan et al. 2009). However, discrete models often lack macroscopic intuition and can be computationally intensive, especially with proliferation and death included, which are commonly stochastic and require many realisations to understand the average behaviour. Continuum models commonly include proliferation and death through source/sink terms and may require constitutive equations to close the system (Antman 2005, Basan et al. 2019, Goriely 2017, Levayer 2019, Matamoro-Vidal et al. 2019, Moulton et al. 2013, Recho et al. 2016, Shraiman 2005). In general, continuum models do not make the underlying cell-level processes clear (Geritz et al. 2012). However, continuum models can be less computationally expensive than discrete models and can be analysed with well-established mathematical techniques such as stability analysis (Armstrong et al. 2006) and phase plane analysis (Landman et al. 2005).

We are most interested starting with discrete descriptions of individual cell dynamics and properties and then deriving corresponding continuum models (Bodnar et al. 2005, Fozard et al. 2010, Matsiaka 2018, O’Dea et al. 2012, Penta et al. 2014, Van Meurs et al. 2019, Yereniuk et al. 2019) because this allows us to switch between the two spatial scales and take advantage of both. Further, this approach is very insightful as it can be used to demonstrate conditions when continuum models are valid and when they are not valid. Having a continuum model which is more computationally efficient to solve than the discrete model, and which well-established mathematical techniques can be applied to, is only beneficial if the continuum model accurately represents the underlying discrete behaviour. In this work, we start with the model of mechanical relaxation in heterogeneous epithelial tissues from Murphy et al. (2019) and now incorporate cell-length-dependent proliferation and death mechanisms. This framework allows us to explore mechanical cell competition, which was not previously possible when considering only homogeneous populations (Murray et al. 2009, 2010, 2011, 2012, Baker et al. 2019) or two populations without cell death (Lorenzi et al. 2019).

This work is structured as follows. In Section 2, we present a new discrete mechanical model that includes cell-length-dependent proliferation and death mechanisms. We then derive the corresponding novel continuum model that takes the form of a system of coupled nonlinear partial differential equations with both hyperbolic and parabolic properties. In Section 3.1, we explore our novel model by considering the evolution of a homogeneous tissue where cells are undergoing both proliferation and death. In Section 3.2, we explore mechanical cell competition in the context of cancer invasion by considering a heterogeneous tissue composed of both cancerous and normal cells that compete for space. Using the model we explore whether cancer cells will eventually replace all of the healthy cells or can the cancer cells coexist with the healthy cells? In Section 3.3, we demonstrate the importance of the discrete to continuum approach.

## 2 Model formulation

In this section, we focus on how we stochastically implement cell proliferation and death for heterogeneous cell populations within the discrete mechanical framework and the derivation of the continuum description.

### 2.1 Discrete model

We start by describing the mechanical model and then include proliferation and death. We represent the epithelial tissue as a one-dimensional chain of cells, connected at cell boundaries, in a fixed domain of length *L*. The cells experience cell-cell interaction forces at their cell boundaries, for example cellcell adhesion (Johnston et al. 2013) or compressive stresses (Tse et al. 2012). For a system of *N* cells, cell *i* has left and right cell boundaries at positions 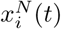, 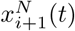, respectively. Fixed boundary conditions at *x* = 0 and *x* = *L* are imposed 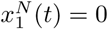 and 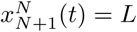. To allow for heterogeneous tissues, each cell *i*, which can be thought of here as a mechanical spring, is prescribed with intrinsic cell properties including a cell stiffness, 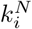, and resting cell length, 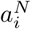 (Figure 1a). We assume cell motion occurs in a viscous environment such that cell boundaries experience a drag force with mobility coefficient *η* > 0 (Fletcher et al. 2014, Matsiaka et al. 2018, Murphy et al. 2019). In the overdamped regime, where inertia effects are neglected, the evolution of cell boundary *i* in a system of *N* cells is

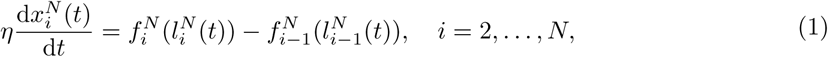

where 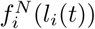 is the force exerted on cell *i* − 1 by cell *i* (Murphy et al. 2019). When 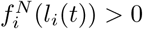 cell *i* contracts and pulls cell *i* − 1. When 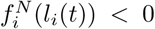 cell *i* extends and pushes cell *i* − 1. This cell-cell interaction force law may be given by, for example, a cubic, Hertz, Lennard-Jones, or Johnson-Kendall-Roberts law (Baker et al. 2019, Lorenzi et al. 2019, Murray et al. 2012). However, for simplicity, we choose a Hookean force law,

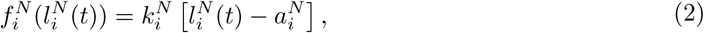

where cell *i* has length 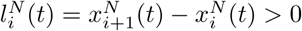.

**Figure 1:**
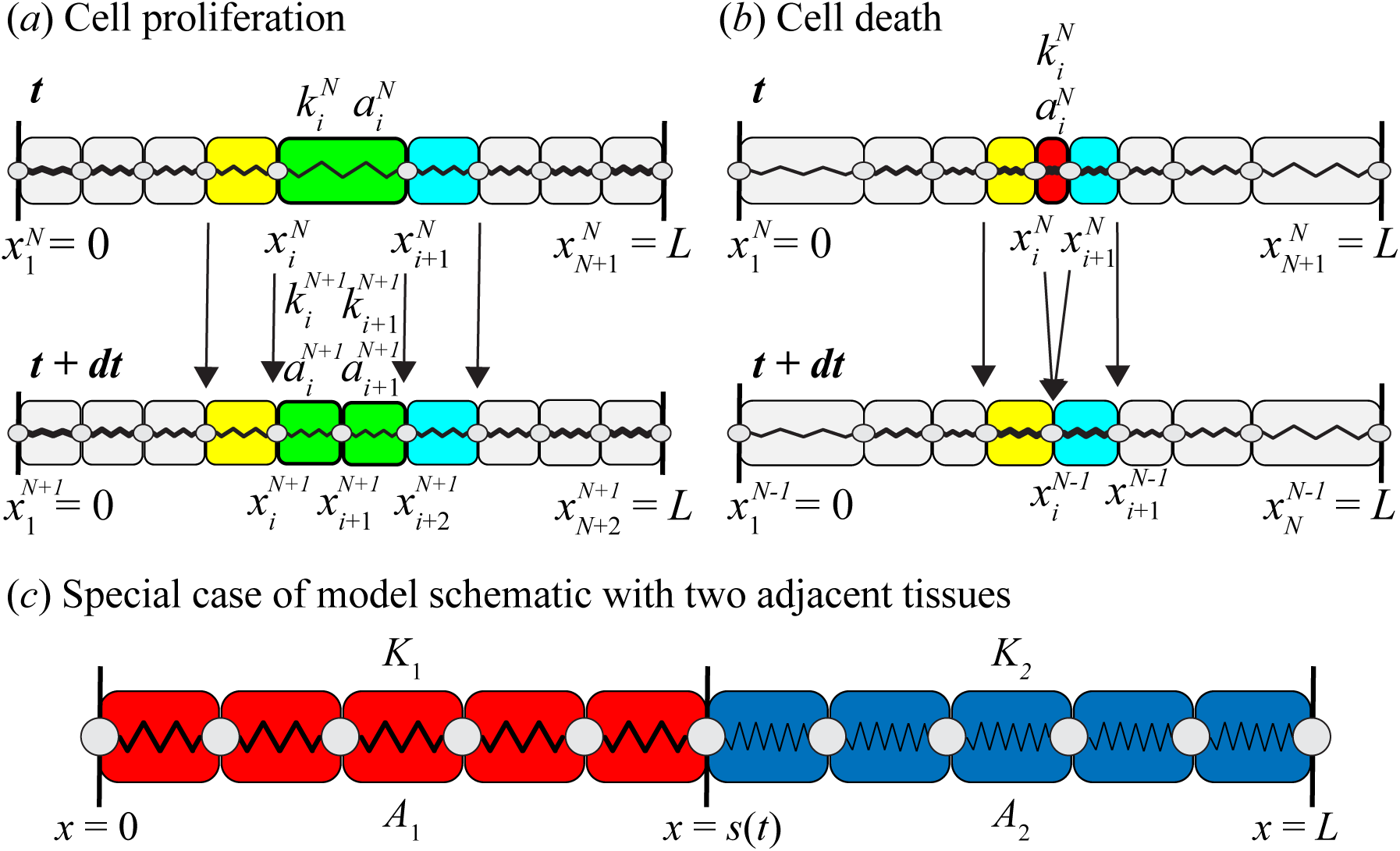
Discrete model schematic for a heterogeneous epithelial tissue with cell proliferation and death. Cell *i* in a system of *N* cells has left and right cell boundaries 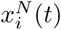, 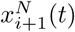, with 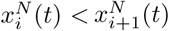, respectively, and is prescribed with a cell stiffness 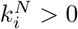, and a resting cell length 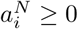. *(a)* Cell proliferation. Cell *i*, coloured green, is selected to proliferate at time *t*. At time *t* + d*t*, the cell has proliferated with a new cell boundary introduced at the midpoint of the original cell. Cell properties of the daughter cell are prescribed from the parent cell. (*b*) Cell death. Cell *i*, coloured red, is selected to die at time *t*. At time *t* + d*t*, the cell is removed and the cell boundaries of cell *i* at time have coalesced at midpoint of the original cell. For both proliferation and death cells are re-indexed at time *t* + d*t*. (*c*) Special case with two adjacent tissues. The left tissue (tissue 1) is coloured red and the right tissue (tissue 2) is coloured blue. The interface position between the left and right tissues is *x* = *s*(*t*). Each cell in tissue *i* has cell stiffness *K*_*i*_ and resting cell length *A*_*i*_. Proliferation and death rates remain dependent on the length of each cell. This could also represent a single tissue with internal heterogeneity.

We include cell proliferation stochastically, by considering that cell *i* proliferates with probability 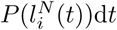 in the small time interval [*t, t* + d*t*), that depends on the current cell length, 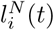, and proliferation mechanism *P* (⋅) (Baker et al. 2019, Puliafito et al. 2012). When cell *i* proliferates we increase the number of cells by one by introducing a new cell boundary, 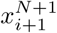, at the midpoint of the original cell, 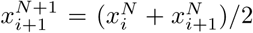, and relabel indices accordingly (Figure 1a). Daughter cells take the same intrinsic cell properties as the parent cell. Cell death is included similarly to cell proliferation with a cell-length-dependent death mechanism, 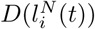. In a system of *N* + 1 cells, when cell *i* dies, with cell boundaries 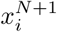 and 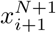, the number of cells is reduced by one. The two cell boundaries are set to instantly coalesce at the midpoint of the dying cell (Figure 1b). Cell death at the tissue boundaries needs to be considered separately (Supplementary Material SM1.2). In this work, we consider constant, linear, and logistic models of proliferation and death (Table 1, Figure 2). We solve discrete Equations (1) together with a stochastic implementation of proliferation and death numerically (Supplementary Material SM2.1).

**Table 1:** Proliferation and death mechanisms written in terms of cell length, 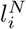, proliferation parameter, *β*, and death parameters, *γ*, *l*_*d*_.

|  | Constant | Linear | Logistic |
| --- | --- | --- | --- |
| $P(l_i^N)$ | $\beta$ | $\beta l_i^N$ | $\beta$ |
| $D(l_i^N)$ | $\gamma$ | $\begin{cases} \gamma(l_d - l_i^N), & 0 \leq l_i^N \leq l_d \\ 0, & l_d < l_i^N \end{cases}$ | $\frac{\gamma}{l_i^N}$ |

**Figure 2:**
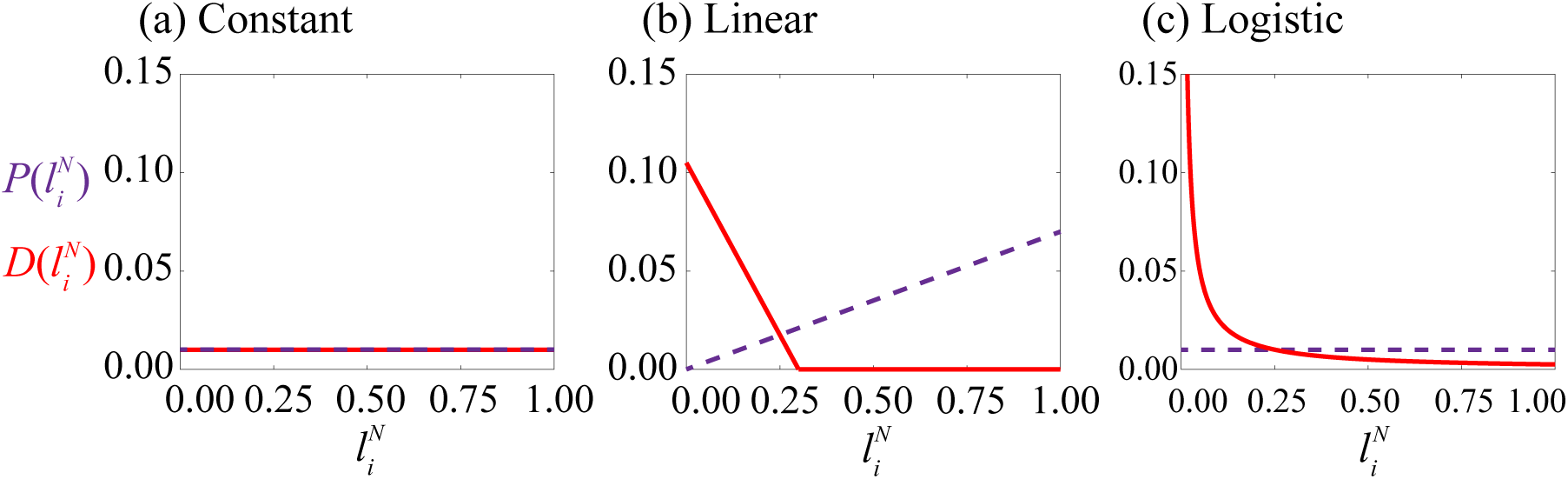
Proliferation and death mechanisms considered in this work. Proliferation rates, 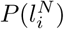 (dashed), and death rates, 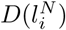 (solid), shown as a function of cell length, 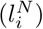. Parameters used in this work: (a) *β* = 0.01, *γ* = 0.01, (b) *β* = 0.07, *γ* = 0.35, *l*_*d*_ = 0.3, (c) *β* = 0.01, *γ* = 0.0025.

### 2.2 Derivation of continuum model

To understand the mean behaviour of the discrete model we must average over many identically prepared stochastic realisations. However, this can be computationally intensive, especially for large *N*. The corresponding continuum model, which we first present and then derive, represents the average behaviour and unlike the discrete model the computational time required to solve the continuum model is independent of *N*.

The continuum model for the evolution of the cell density, *q*(*x, t*), in terms of the continuous cell-cell interaction force, *f* (*x, t*), proliferation rate, *P* (1*/q*(*x, t*)), and death rate, *D*(1*/q*(*x, t*)), is the conservation of mass equation

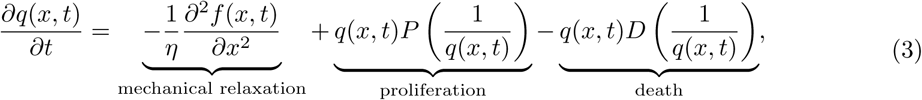

where the continuous cell-cell interaction force which corresponds to Equation (2) is given by

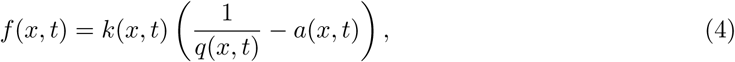

with cell stiffness, *k*(*x, t*), and the resting cell length, *a*(*x, t*), also being described by continuous fields. From Equation (3), we know that the cell density flux, *j*(*x, t*) = *q*(*x, t*)*u*(*x, t*), is equal to the spatial gradient of the cell-cell interaction force, (1*/η*)∂*f/∂x*. Therefore, the cell velocity, *u*(*x, t*), is related to the cell density and gradient of the cell-cell interaction force through

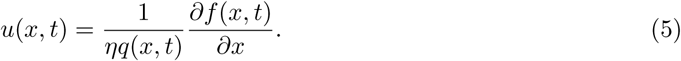

Note that Equation (5) corresponds to the discrete linear momentum equation in Equation (2). Intrinsic mechanical cell properties are constant for each cell and transported by the motion of cells. The proliferation and death functions, *P* (⋅) and *D*(⋅), respectively, (Table 1) are evaluated at 1*/q*(*x, t*). Depending on the choice of proliferation and death mechanisms we may have additional intrinsic cellular properties, β(*x, t*)*, γ*(*x, t*), and *l*_*d*_(*x, t*). All intrinsic cellular properties evolve according to the following transport equation,

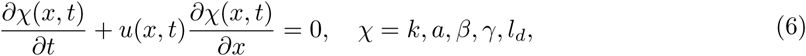

where *u*(*x, t*) is the cell velocity. The left hand side of Equation (6) corresponds to the material derivative, expressing the fact that there is no change in cellular properties along cell trajectories. We solve the system of Equations (3)–(6) together with initial conditions and boundary conditions numerically (Supplementary Material SM2.2).

We now systematically derive Equation (3). We take care to explicitly state and make clear all approximations made in this derivation. We incorporate proliferation and then death into the modelling framework, under the assumption that the two processes are independent. The previously derived mechanical relaxation term and transport of cellular property equations (6) are briefly discussed (Murphy et al. 2019). For clarity, the derivation is shown for one spring per cell. However, this analysis can be extended to *m* > 1 springs per cell which, for sufficiently small *N*, is a more appropriate method to define the continuous field functions (Murphy et al. 2019) (Supplementary Material SM1.3).

#### 2.2.1 Proliferation

As cell proliferation is included stochastically (Sections 2.1, SM2.1), we consider an infinitesimal time interval [*t, t* + d*t*) and condition on the possible proliferation events that could occur and influence the position of cell boundary *i* in a system of *N* cells. Choosing d*t* sufficiently small so that at most one proliferation event can occur in [*t, t* + d*t*), there are four possibilities: i) there is no proliferation, in which case the cell boundary position 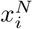 only changes by mechanical relaxation; ii) there is proliferation to the right of cell *i* − 1; iii) there is proliferation to the left of cell *i* − 1; and iv) cell *i* − 1 proliferates. This leads to the following infinitesimal evolution law for the position of cell boundary 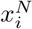, accounting for cell relabelling when a new cell is added:

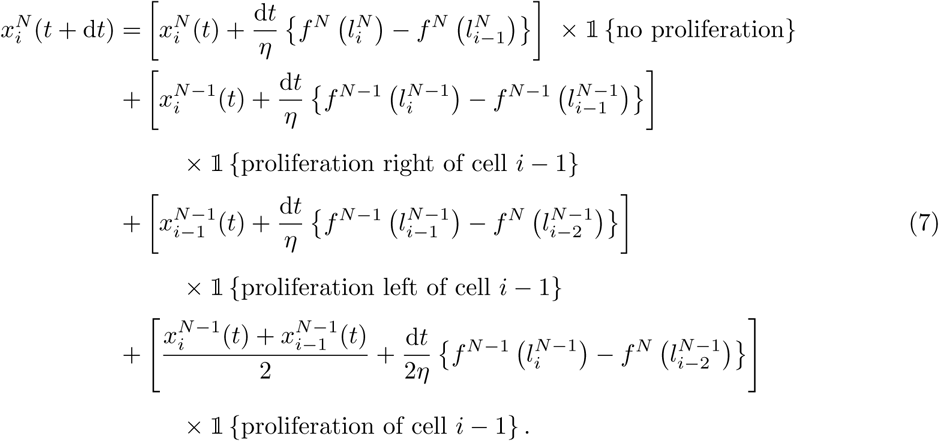

Each term in square brackets is the resulting force from neighbouring cells due to mechanical relaxation, given by Equations (1), for each potential event. In addition, we include Boolean random variables expressed as indicator functions, 𝟙 *{⋅}*, defined as

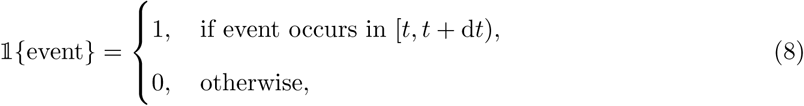

whose expectations in the context of Equation (7) can be interpreted as proliferation probabilities. For a system of *N* cells, where d*t* is sufficiently small, these proliferation probabilities are given by

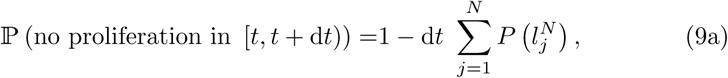

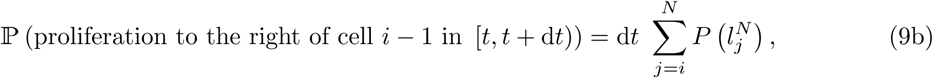

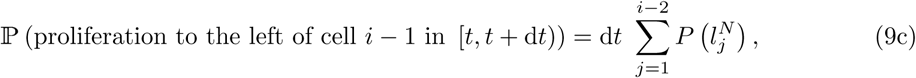

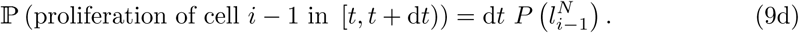

Taking a statistical expectation, denoted *〈⋅〉*, of Equation (7), 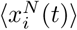 now represents the expected position of cell boundary *i* at time *t* in a system of *N* cells. We use the proliferation probabilities with the following simplifying assumptions: i) 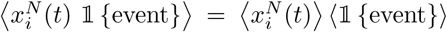, namely independence of the position of the cell boundary in space and proliferation propensity, and a meanfield approximation as proliferation propensities depend on cell length; ii) 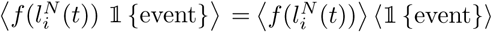, namely independence of the force and the propensity to proliferate, and a mean-field approximation as force depends on cell length; iii) a statistical mean-field approximation for force, 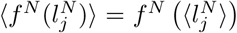, and proliferation propensities, 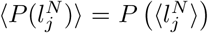. For simplicity we now drop the *〈⋅〉* notation. Then,

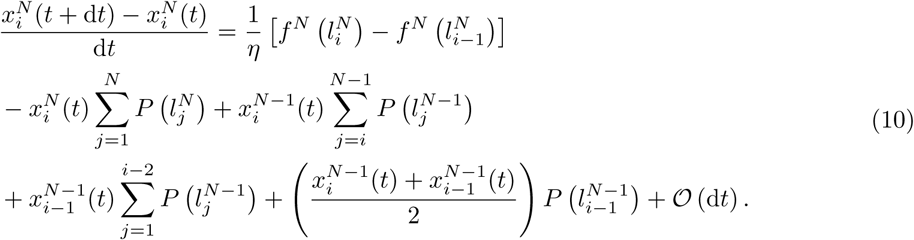

We also assume: iv) the total propensity to proliferate is not significantly changed due to single a proliferation event, 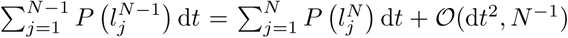; v) a single proliferation event does not significantly alter the position of a cell boundary, 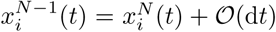 (Figure 1). As we will show, assumptions iv) and v) are good approximations for large *N* and allow us to combine summations. Then, assuming vi) 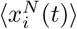 is a continuous function of time, we rearrange and take the limit d*t* → 0. For the proliferation terms we replace the cell length with the discrete cell density 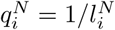 to obtain

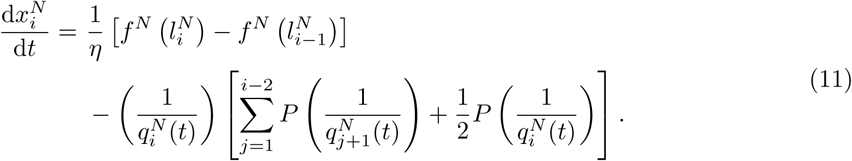

Equation (11) is only valid for the time interval [*t, t* + d*t*) under the assumptions iv) and v) above. Thus far, we have extended the discrete model with mechanical relaxation to include the effects of cell proliferation. However, the statistically averaged model still retains information about discrete cell entities. We thus average over space to define a continuum cell density. Following Murphy et al. (2019), we introduce the microscopic density of cells,

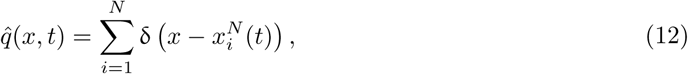

where δ is the Dirac delta function (Evans et al. 2008, Lighthill 1958). We define a local spatial average over a length scale δ*x*, denoted (⋅)δ_*x*_, such that *a*_*i*_ ≪ δ*x* ≪ *L*, which is sufficiently large to capture local heterogeneities for cellular properties that are constant during cell motion, including *k* and *a*, but sufficiently small to define continuous properties across *L*. The continuous cell density function, *q*(*x, t*), is thus defined as

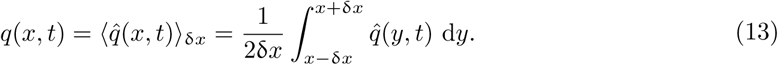

Differentiating Equation (13) with respect to time gives

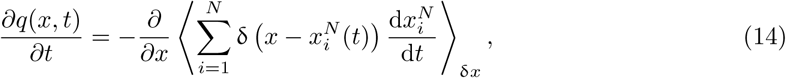

where we use properties of the Dirac delta distribution (Lighthill 1958) and interchange the derivative with the spatial average as δ*x* is small. Consistent with assumptions iv)-v) above, the sum over the microscopic densities can be considered to be fixed over *N* cells in Equation (14) within the small time interval [*t, t* + d*t*).

On the right hand side of Equation (11), the first two terms involving *f* correspond to a mechanical contribution. This contribution is unchanged compared to Murphy et al. (2019) and, when substituted into Equation (14), it gives rise to the mechanical relaxation term in the right hand side of continuum model Equation (3) (Supplementary Material SM1.4). We now focus only on the contribution due to proliferation determined by substituting the proliferation terms of Equation (11) into Equation (14), giving a contribution which we denote ∂*q*(*x, t*)*/∂t*|_*P*_,

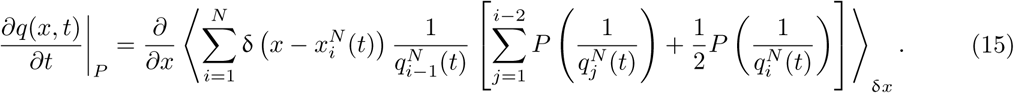

Now, assuming vii) that the spatial average interval is sufficiently far from the tissue boundary, i.e. *i* ≫ 1, we make the following approximation:

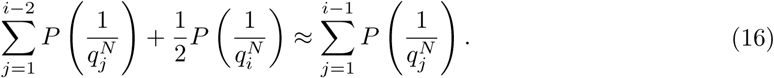

To switch the dependence on the cell index to cell position, we multiply each term indexed by *j* in the sum on the right hand side of Equation (16) by 1 = *l*_*j*_*q*_*j*_. Then, relating the discrete cell density to the continuous density through 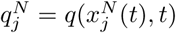, gives

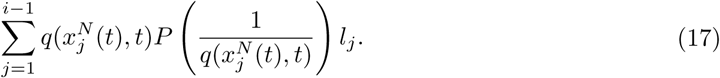

We discretise the spatial domain *x*_1_ ≤ *x* ≤ *x*_*i*−1_ with a uniform mesh with nodes *y*_*s*_, *s* = 1, 2, …, *S*, where *y*_1_ = *x*_1_, *y*_*S*_ = *x*_*i*−1_, and *y*_*s*_ − *y*_*s*−1_ = ∆ *y* ≪ *l*_*j*_. Then, evaluating the continuous density at each node position, *y*_*s*_, we interpret Equation (17) as the following Riemann sum

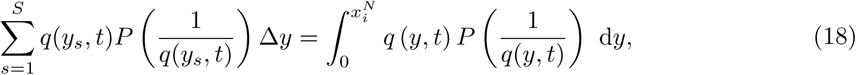

where the integral on the right hand side is obtained by taking the limit ∆*y* → 0. Substituting Equation (18) into Equation (15) gives

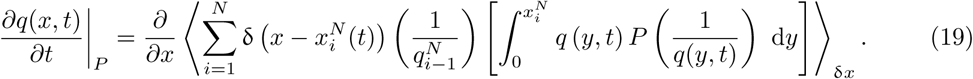

Calculating the spatial average, which only includes contributions from within the spatial average interval due to the Dirac delta functions, gives

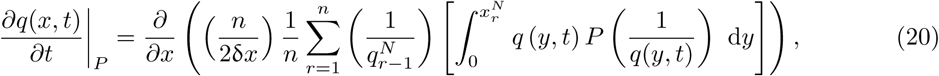

where the index *r* labels the *n* cell boundaries contained within the spatial average interval (*x* − δ*x, x* + δ*x*). Since *a*_*i*_ ≪ δ*x* ≪ *L* and *n* ≫ 1 we have 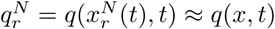 for all *r*, which is now independent of *r*. Similarly, 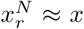 for all *r*, where *x* is the centre of the spatial average interval. This gives

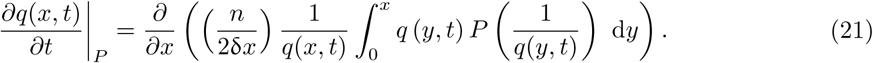

As *n*/(2δ*x*) = *q*(*x,t*) in this spatial average interval, Equation (21) simplifies to

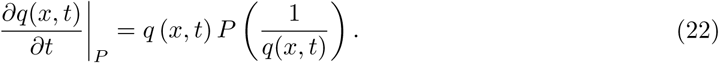

At this point, we see that all explicit references to the total number of cells, *N* (*t*), vanish. This allows the validity of the derivation, initially restricted to the time interval [*t, t* + d*t*), to be extended to arbitrary times. As 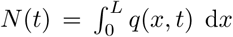, the change in the total cell number with time due to proliferation is accounted for through the source term written in Equation (22). We also stated assumption vii) that held true when sufficiently far from the tissue boundary but we find that this works at the boundary also (Sections 3.1, 3.2). Equation (22) shows proliferation arises as a single source term consistent with usual continuum-based formulations of proliferation whereas in Baker et al. (2019) proliferation arises as this term with an additional contribution.

#### 2.2.2 Death

The derivation of the cell death sink term follows similarly to that of the cell proliferation source term. We again consider an infinitesimally small time interval [*t, t* + d*t*), so that at most one cell death event can occur in [*t, t* + d*t*), and condition on cell death events to understand all possible events that occur and influence cell boundary *i* at *t* + d*t*. This gives

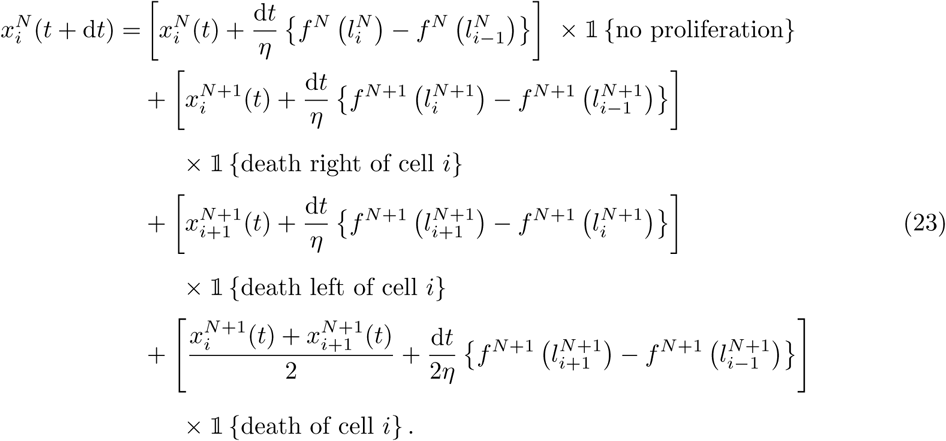

The cell death probabilities for Equation (23) for a system of *N* cells are given by

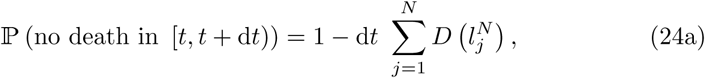

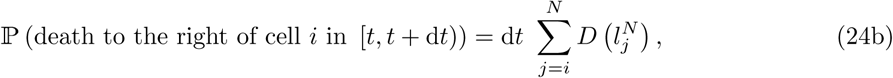

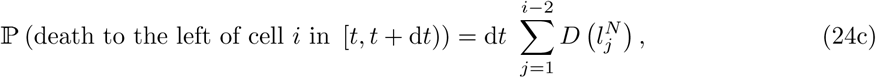

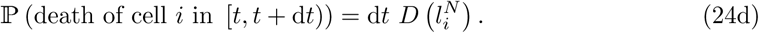

Proceeding similarly to the proliferation derivation, we obtain

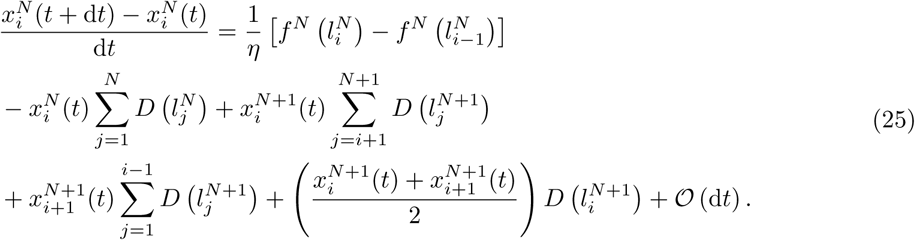

Then, following the same approach as the proliferation derivation, we arrive at the sink term in Equation (3) for cell death, −*q*(*x, t*)*D*(1*/q*(*x, t*)).

#### 2.2.3 Cell properties

Each cell is prescribed with intrinsic mechanical, proliferation, and death properties which are taken to be constant for each cell throughout the simulation. For mechanical cell properties, which include cell stiffness and resting cell length, we have the relationships 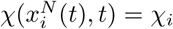 for *χ* = *k, a*. Similar relationships can be defined for the proliferation and death cell properties, *β*, *γ*, *l*_*d*_. Differentiating these equations with respect to time we obtain Equations (6) (Murphy et al. 2019).

## 3 Numerical results

In this section, we first explore the evolution of a homogeneous tissue with different proliferation and death mechanisms and then explore mechanical cell competition for a heterogeneous tissue. To conclude we demonstrate the importance of the discrete to continuum approach through a series of problems where we compare the averaged discrete data with solutions of the corresponding continuum equations.

### 3.1 A homogeneous tissue

The simplest case to consider first is a homogeneous tissue composed of a population of identical cells. We explore three different proliferation and death mechanisms: constant, linear, and logistic (Table 1, Figure 2). For each mechanism we explore proliferation only, death only, and proliferation with death. Cell proliferation and death parameters (Figure 2) are chosen to observe homeostasis where the total cell number remains stable at approximately *N* (*t*) = 40 for *t* > 0.

In all simulations we set *L* = 10, use a Gaussian initial density centred at *x* = *L*/2 with variance three and scaled to have *N* (0) = 40. We set *k* = 10, so that mechanical relaxation is fast in comparison to the proliferation and death (Baker et al. 2019). For individual realisations this results in uniform densities except for short-time transient behaviour following a cell proliferation or death event (Figures 3a-c, 4a-c, S4a-c). Since epithelial cells in a tissue are in extension (Wyatt et al. 2016), we set *a* = 0 for simplicity.

**Figure 3:**
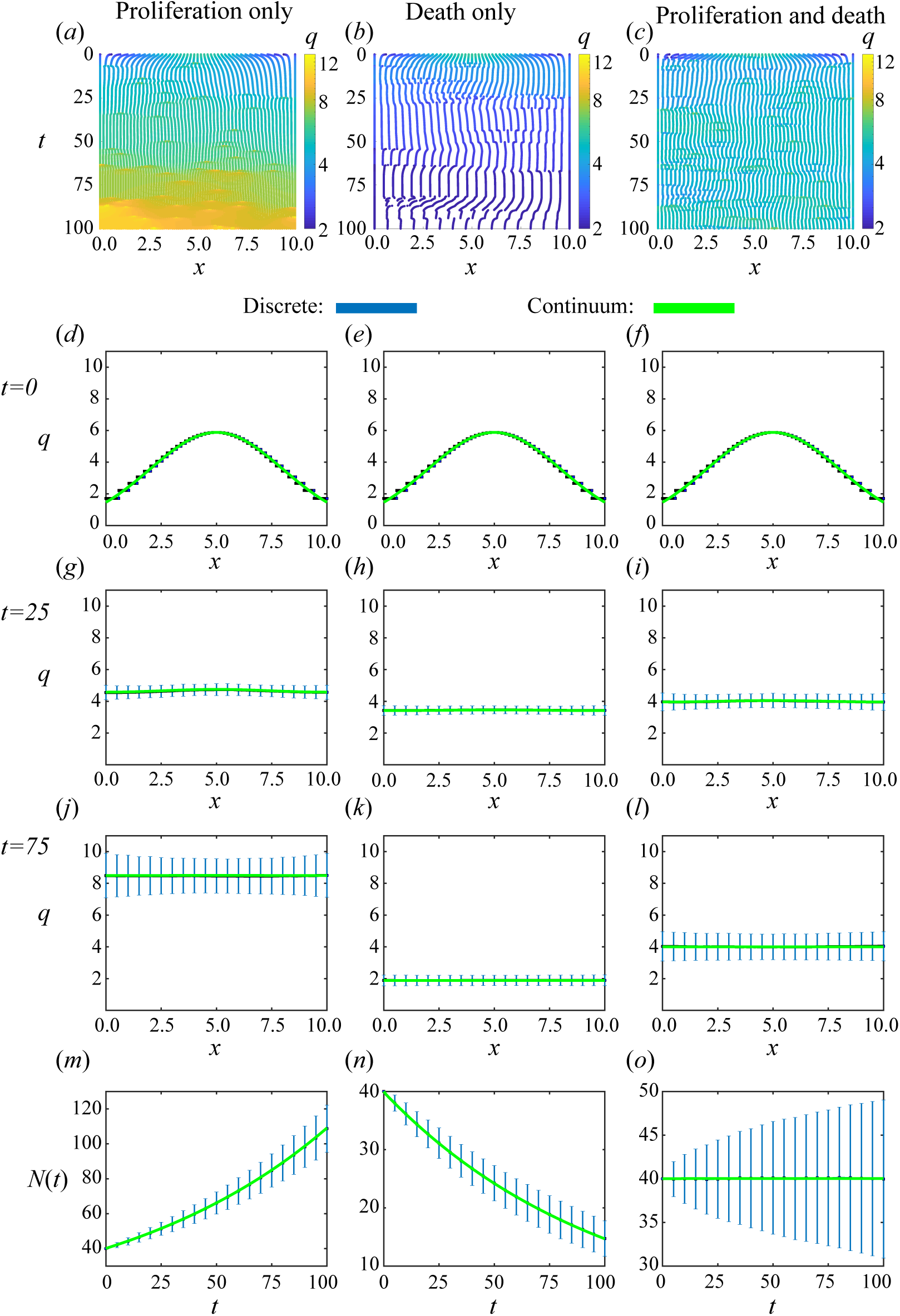
Homogeneous population with **constant** proliferation and death mechanisms. Proliferation only, death only, and proliferation with death shown in the left, middle and right columns, respectively. (a)-(c) Single realisations of cell boundary characteristics for 0 ≤ *t* ≤ 100. (d)-(f), (g)-(i), (j)-(l) Density snapshots at times *t* = 0, 25, 75, respectively. (m)-(o) Total cell number. The average and standard deviation (blue error bars) of 2000 discrete simulations are compared to solution of continuum model (green).

For individual discrete realisations, cell proliferation causes a localised force imbalance followed by fast mechanical relaxation towards mechanical equilibrium and an overall increase in density (Figures 3a, 4a,S5a). Similarly, cell death results in a decrease in density followed by fast mechanical relaxation and an overall decrease in density (Figures 3b, 4b, S5b). With proliferation and death, cell boundaries are repeatedly introduced and removed, and the overall density remains, on average, constant (Figures 3c, 4c, S5c).

We observe excellent agreement when we compare the mean of many identically prepared discrete realisations and the corresponding solutions of the continuum model for both density snapshots and total cell number (Figures 3d-o, 4d-o, S5d-o).

We note that the continuum model does not always provide a good match with an individual realization of the discrete model. For example, for constant proliferation and constant death with equal rates, every discrete realization will eventually become extinct (Supplementary Material SM3.1) as proliferation and death are independent of mechanical relaxation. This is expected as the total cell number is a linear birth-death process (Ross 1996) where the net proliferation rate is always equal to zero (Figure S4). As a consequence, the standard deviation of the total cell number increases with time (Figures 3o). When cell proliferation and death are cell-length-dependent there is closer agreement between the continuum model and single realisations. The net proliferation rate adjusts, due to changes in the number of cells and their lengths, to stabilise the population at its equilibrium value (Figure S4). Therefore extinction is extremely unlikely and the standard deviation of averaged discrete realisations is smaller (Figures 4o, S5o).

**Figure 4:**
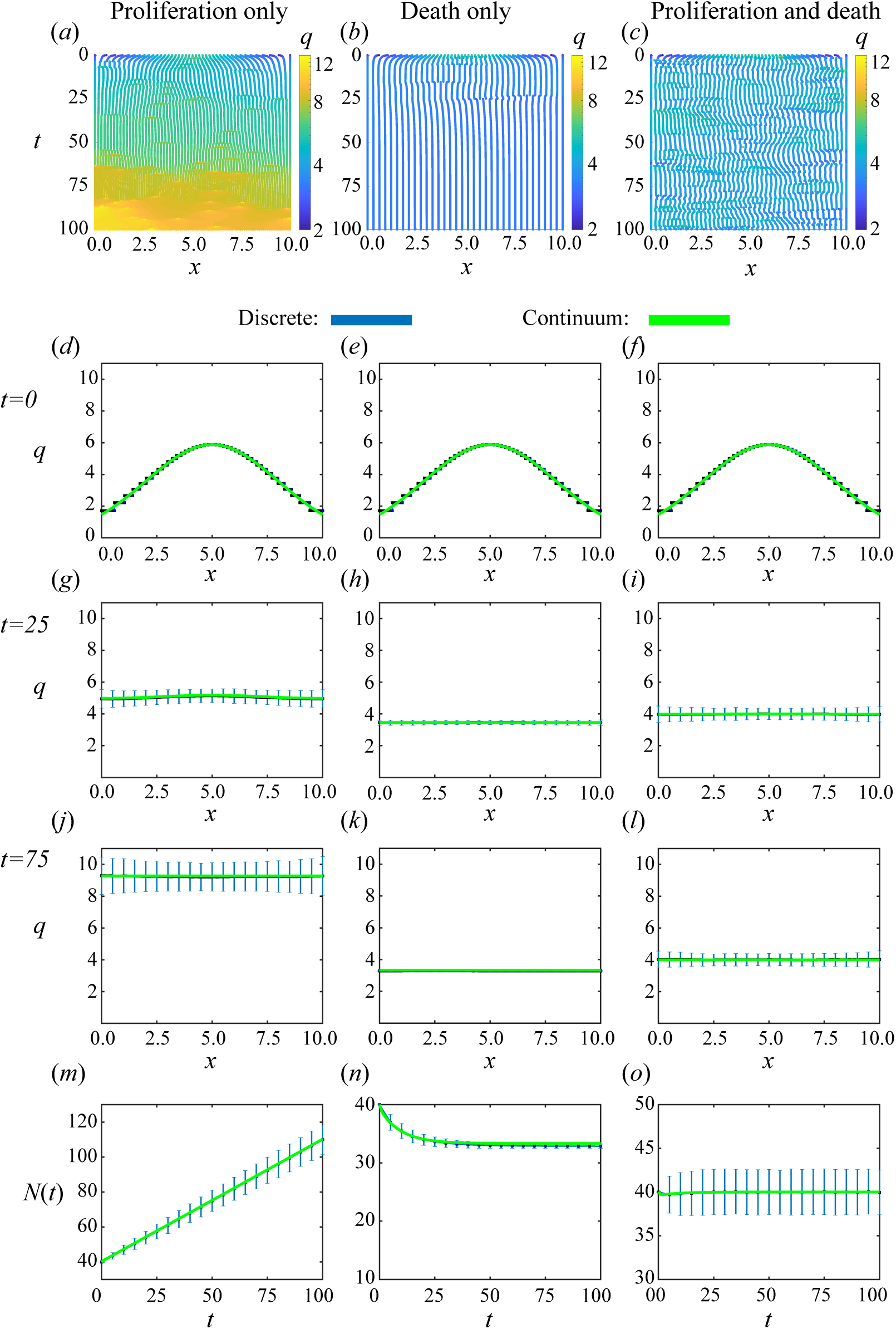
Homogeneous population with **linear** proliferation and death mechanisms. Proliferation only, death only, and proliferation with death shown in the left, middle and right columns, respectively. (a)-(c) Single realisations of cell boundary characteristics for 0 ≤ *t* ≤ 100. (d)-(f), (g)-(i), (j)-(l) Density snapshots at times *t* = 0, 25, 75, respectively. (m)-(o) Total cell number. The average and standard deviation (blue error bars) of 2000 discrete simulations are compared to solution of continuum model (green).

### 3.2 Mechanical cell competition

How tissues compete with each other for space is of great interest with many open biological questions being pursued in the experimental cell biology literature (Bras-Pereira et al. 2018, Levayer 2019, Tsuboi et al. 2018). For example, in cancer invasion in an epithelial tissue a key question is whether cancer cells will eventually replace the entire healthy tissue or can the cancer cells coexist with the healthy cells? We consider this question by simulating a heterogeneous tissue composed of two populations, cancer cells adjacent to healthy cells (Figure 1c). Biologically, it is a hallmark of cancer cells that they are more proliferative and resistant to death than healthy cells (Hanahan et al. 2011). In existing models the standard procedure would be to include these hallmarks as modelling assumptions and not consider the role of mechanical relaxation. However, we will now show this assumption is not necessary. We find that mechanical differences are sufficient for these hallmarks to arise and for cancer cells to outcompete healthy cells. We prescribe cancer and healthy cells the same proliferation and death mechanisms and parameters. We ask a further key question, how does mechanical relaxation alone compare to mechanical relaxation with proliferation, and to mechanical relaxation with proliferation and death?

In all scenarios, the left tissue (tissue 1) is coloured red to represent cancer cells and the right tissue (tissue 2) is coloured blue to represent healthy cells (Figure 1c). Each tissue starts with 20 cells. We assume cancer cells have lower stiffness than healthy cells (Lekka 2016) so we set cells in tissue 1 and 2 with cell stiffnesses *K*_1_ = 10 and *K*_2_ = 20, respectively. Again, for simplicity and to represent that cells in an epithelial tissue are understood to be in extension (Wyatt et al. 2016), we set *a* = 0.

With only mechanical relaxation the interface position, *s*(*t*), relaxes to the long-time interface position, 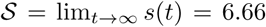 (Murphy et al. 2019). In this scenario, the cancer and healthy cells coexist. However, the assumption of mechanical relaxation alone is only realistic over a short timescale where proliferation and death are negligible. When we include proliferation and death below, we use this long-time solution as the initial condition. As the mechanical relaxation rate is faster than the proliferation and death rates, using this initial condition only neglects initial short-time transient behaviour and does not significantly impact the long-time solution.

For mechanical relaxation with proliferation (Figure 5), we prescribe the linear proliferation mechanism for both the cancer and healthy cells with the same parameters. As cancer cells have lower cell stiffness than healthy cells, the cancer cells are always longer than the healthy cells (Supplementary Material SM4.1) except for the short transients after proliferation events where the cells have yet to mechanically relax. Initially, the cancer cells, with length 1/3, are double the length of healthy cells. Referring to Figure 2b we see that the difference in cell lengths corresponds to cancer cells being more likely to proliferate than the healthy cells. Therefore, the cancer cells proliferate more than the healthy cells not because they were set to have advantageous intrinsic proliferation or death properties through a modelling assumption, but simply due to the coupling of mechanical relaxation with the length-dependent proliferation mechanism. With each proliferation event all cells become smaller, with the healthy cells remaining smaller than the cancer cells. Here we have coexistence but all cells will eventually become unrealistically small and this happens first for healthy cells. In the absence of cell death, changing the proliferation mechanism will still result in coexistence.

**Figure 5:**
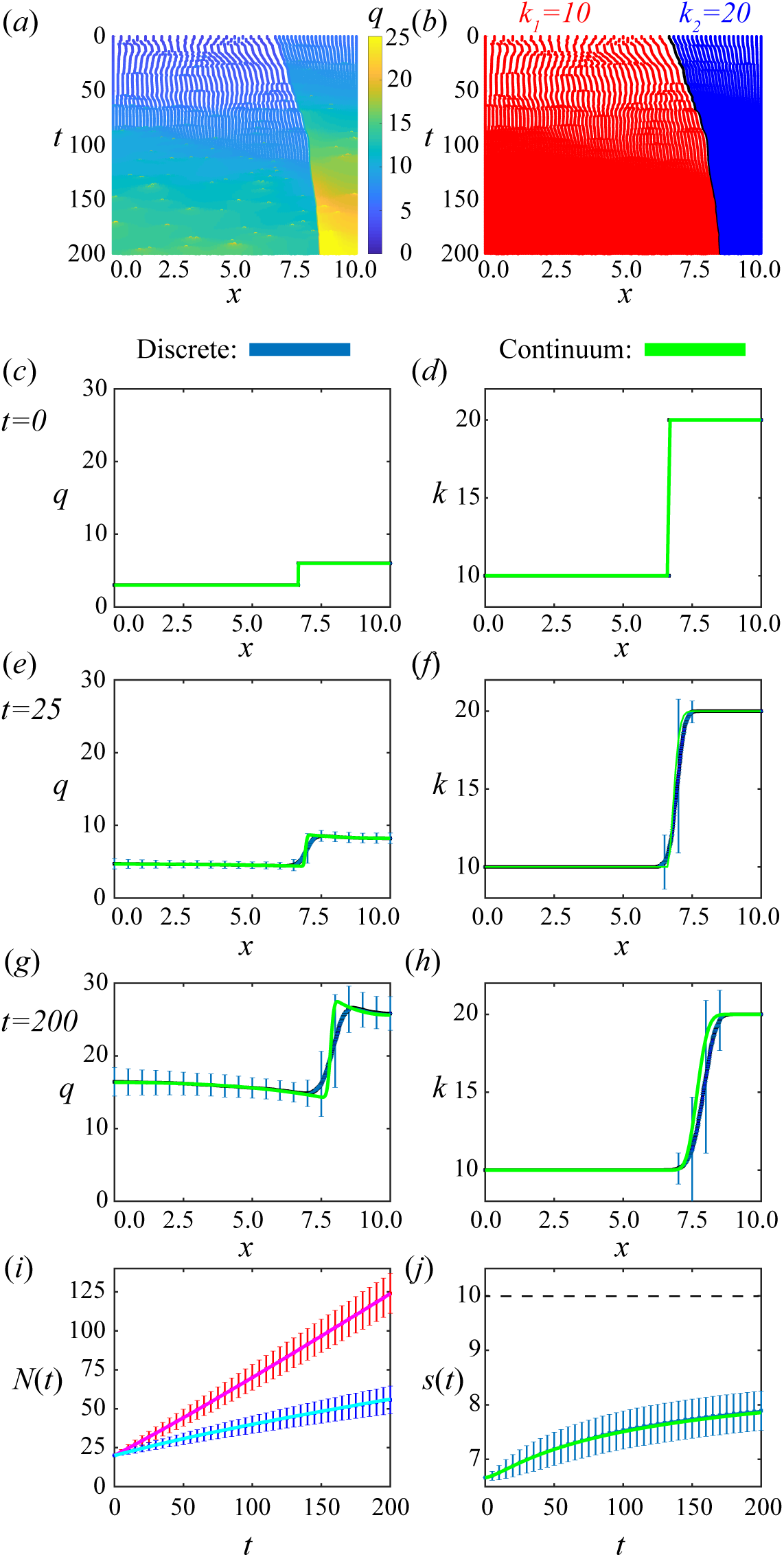
Results for cancer invasion with adjacent populations using **linear** proliferation and death mechanisms with **proliferation only**. (a),(b) A single realization of cell boundary characteristics for 0 ≤ *t ≤* 200. Colouring in (a),(b) represents cell density and cell stiffness, respectively. (c)-(d), (e)-(f), (g)-(h) Density and cell stiffness snapshots, left and right, respectively, at times *t* = 0, 25, 200, respectively. (i) Total cell number, *N*(*t*) > 0, for cancer (red/magenta) and healthy cells (blue/cyan) for the discrete/continuum solutions. (j) Interface position, *s*(*t*), where the dotted line shows the edge of the domain. The average and standard deviation (blue error bar) of 2000 discrete simulations are compared to the solution of the continuum model (green).

For mechanical relaxation with proliferation and death (Figure 6) a cell is more likely to die when it is smaller (Figure 2b). As we have observed for mechanical relaxation with proliferation, the healthy cells are smaller first, due to their higher relative stiffness, and therefore are more likely to die first. Once all of the healthy cells have died we have a homogeneous population of cancerous cells (Section 3.1). Importantly, we find that the cancerous cells, despite having identical proliferation and death mechanisms, are the *winner* cells of mechanical cell competition; they outcompete the healthy cells and take over the domain purely as a result of having lower cell stiffness.

**Figure 6:**
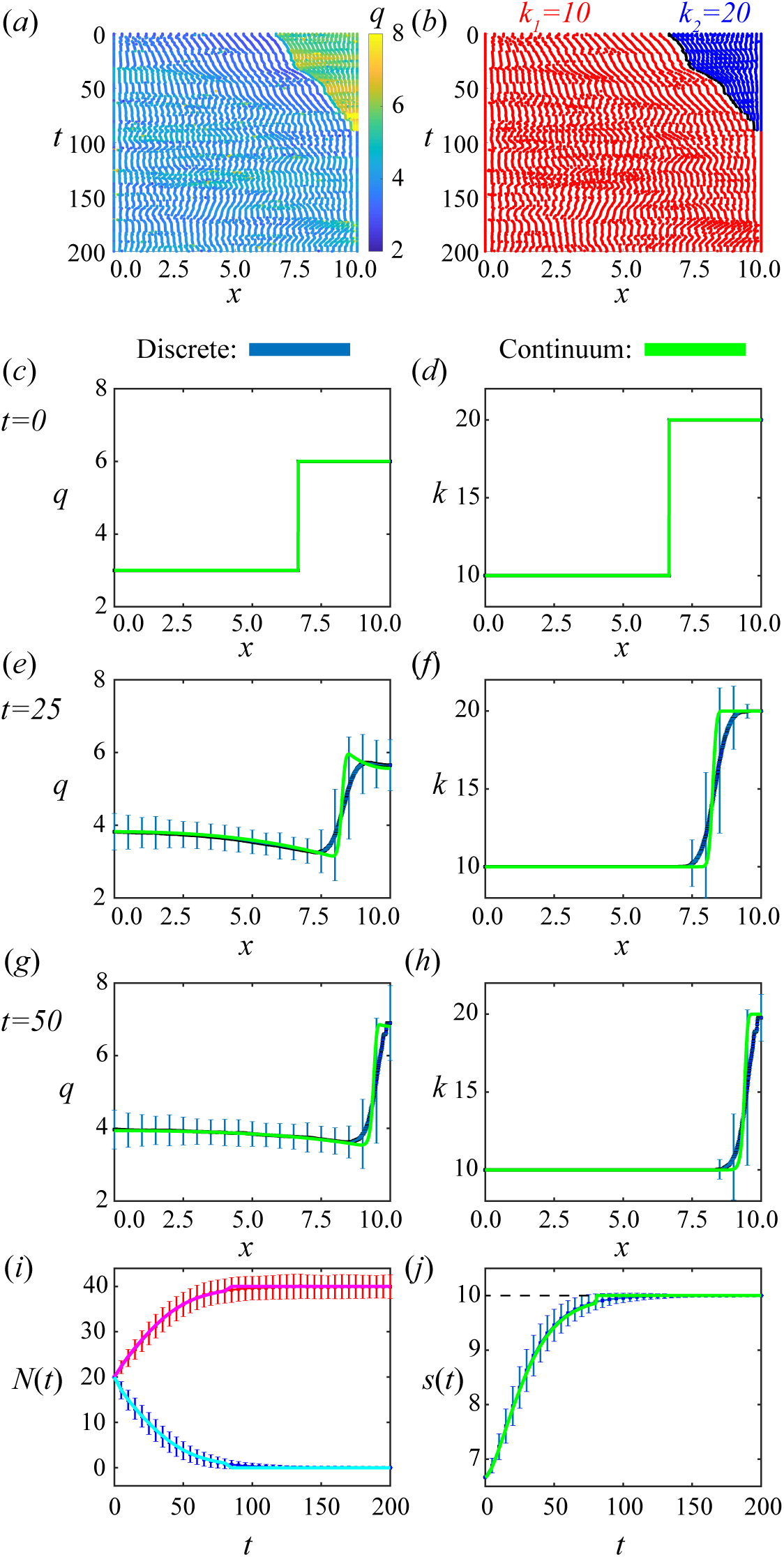
Results for cancer invasion with adjacent populations using **linear** proliferation and death mechanisms with **proliferation and death**. First row shows a single realization of cell boundary characteristics for 0 ≤ *t* ≤ 200. Colouring in (a),(b) represent cell density and cell stiffness, respectively. (c)-(d), (e)-(f), (g)-(h) Density and cell stiffness snapshots, left and right, respectively, at times *t* = 0, 25, 50, respectively. (i) Total cell number, *N* (*t*) > 0, for cancer (red/magenta) and healthy cells (blue/cyan) for the discrete/continuum solutions. (j) Interface position, *s*(*t*), where the dotted line shows the edge of the domain. The average and standard deviation (blue error bar) of 2000 discrete simulations are compared to the solution of the continuum model (green).

Similar results regarding cancer invasion are found when considering the logistic mechanisms with both proliferation and death (Supplementary Material SM4.2). In contrast, for the constant proliferation and death mechanisms, where the proliferation and death mechanisms are both independent of the cell length and therefore independent of mechanical relaxation, to observe cancer cells invading the full domain we would have to prescribe the cancer cells to be more proliferative and resistant to death than the healthy cells.

### 3.3 Importance of the discrete to continuum approach

The discrete to continuum approach is important as it provides a principled means to determine how cell-level properties scale to the macroscale. Further, the approach provides conditions for whether or not the continuum model is beneficial, as we now explore.

In previous sections we choose proliferation and death mechanisms with parameters which lead to a good match between the appropriately averaged data from repeated discrete realisations and the solution of the corresponding continuum model. In Section 3.1, we demonstrate that individual realisations of the discrete model can go extinct while the continuum model does not. This provides a first indication that the continuum model does not always capture all relevant information from the underlying discrete model. We now demonstrate that if the approximations outlined in the derivation of the continuum model in Section 2.2.1 are not satisfied then the continuum approximation is not always satisfactory, and in such cases the discrete model should be used.

As an illustrative example we consider a proliferation mechanism which varies rapidly with cell length. For simplicity we consider the following piecewise cell-length-dependent proliferation mechanism

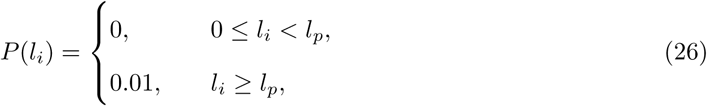

where we set the proliferation threshold to be *l*_*p*_ = 0.2. As before, *N* (0) = 40 but we now choose a constant initial density condition so *l*_*i*_ = 0.25 for each cell. Therefore, in the discrete model, each cell is initially able to proliferate. When the first cell proliferates it divides into two equally sized daughter cells with lengths *l*_*i*_ = 0.125. With fast mechanical relaxation, i.e. sufficiently large *k*, all 41 cells relax to equal size, *l*_*i*_ = 0.244, before the next proliferation event. This repeats until *l*_*i*_ < *l*_*p*_ for each cell *i* when proliferation stops (Figure 7d,f,h). This results in a tissue with 50 cells (Figure 7i,j), which is consistent with the continuum model where the density increases at the same rate everywhere in the tissue until reaching *N* = 50 (Figure 7j, S6b,d,f). As the initial density condition is uniform the continuum solution holds true for any *k*. However, the behaviour of the discrete model for very slow mechanical relaxation, i.e. sufficiently small *k*, is very different. Proliferation occurs faster than mechanical relaxation so each of the initial 40 cells can proliferate, resulting in 80 cells (Figure 7c,e,g,i, S6a,c,e). It is clear that the continuum model does not accurately describe this problem and so we conclude that the discrete model should be used in this case. Increasing *k* results in an improved agreement between the discrete and continuum models (Figure S7).

This example is important. The mismatch between the continuum and discrete results for this case remains even if we consider similar problems with larger numbers of cells, so simply increasing *N* (0) does not alleviate the issue. We do observe that increasing the mechanical relaxation rate, by increasing *k*, does provide a better match. However, in this piecewise proliferation mechanism example we require very high values of *k*, for example *k* = 1000, for a good match. Results in previous sections with excellent agreement are generated using *k* = 10. Revisiting the mechanical cell competition example and reducing to *k* = 0.0001 still provides a reasonable match (Figure S10). This is because the rates involved in the proliferation and death mechanisms are smoother and slowly varying with respect to cell-length.

**Figure 7:**
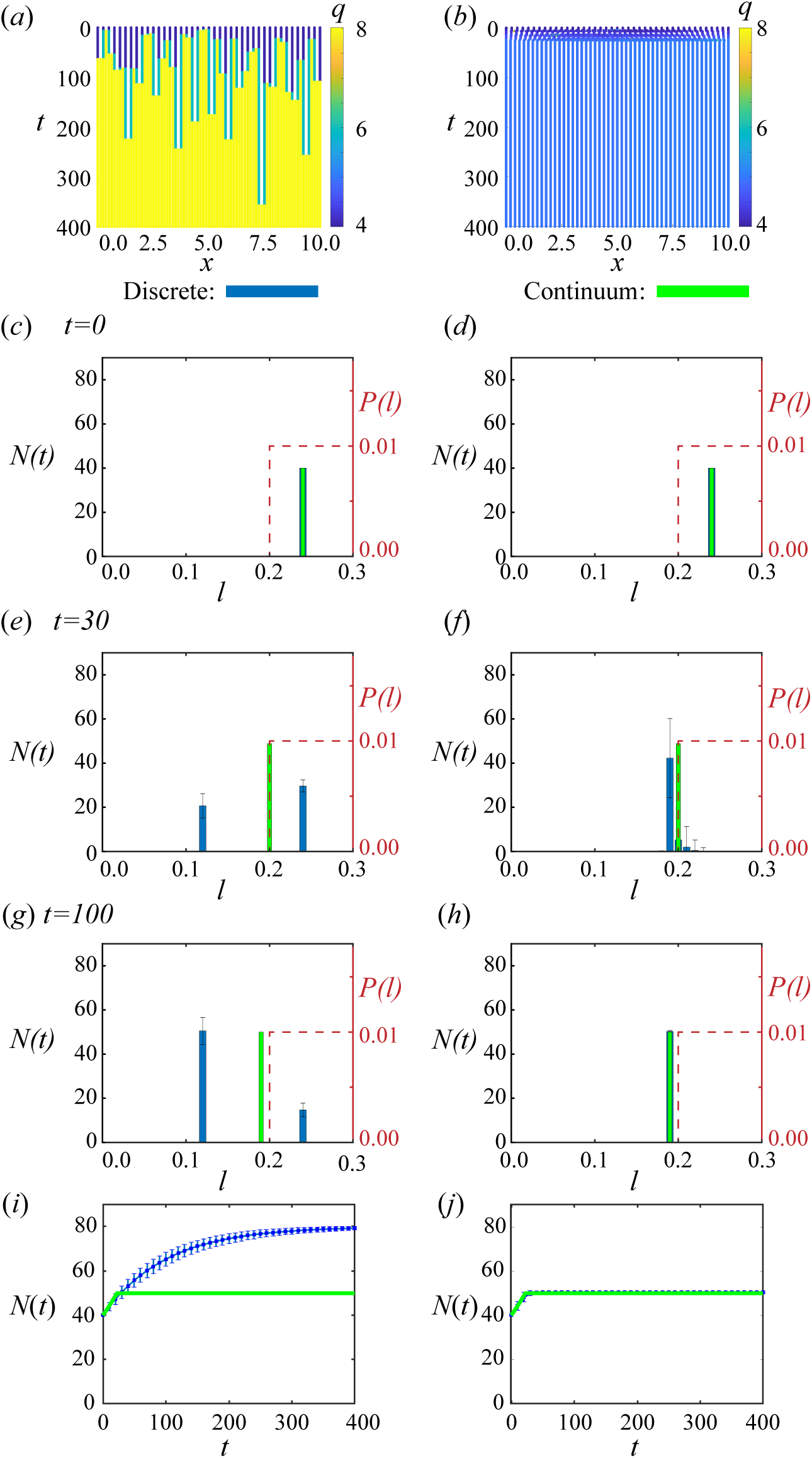
Homogeneous population with rapidly varying proliferation mechanisms. With slow mechanical relaxation, *k* = 0.0001, and faster mechanical relaxation, *k* = 1000, shown in left and right columns, respectively. (a)-(b) Single realisations of cell boundary characteristics for 0 ≤ *t* ≤ 100. (c)-(h) Cell length distributions against proliferation mechanism for times *t* = 0, 50, 100 where one discrete realisation (blue) is compared against continuum model (green). (i)-(j) Total cell number where the average and standard deviation (blue error bars) of 2000 discrete simulations are compared to solution of continuum model (green).

The results in Figure 7 may be surprising from the perspective of continuum mechanics. A common approach in continuum mechanics (Antman 2005, Goriely 2017, Moulton et al. 2013) is to start with conservation of mass and linear momentum and invoke constitutive laws. To derive our model using this approach one could start with the conservation of mass equation and then heuristically add source and sink terms to represent proliferation and cell death to give

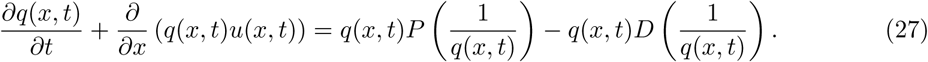

The continuous analogue of the discrete conservation of momentum Equation (1) could be written by expanding the discrete cell-cell interaction force law with respect to cell-length in a Taylor series expansion to obtain

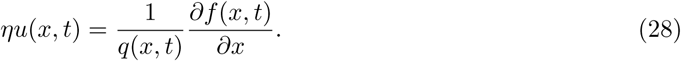

Equations (27) and (28) agree with Equations (3)–(6) derived earlier using a systematic coarsegraining approach. However, in the common continuum mechanics approach we would not have any opportunity to compare solutions of these continuum models with any underlying discrete description. This simple approach does not give any explicit indication of the underlying approximations inherent in the continuum model nor does it inform us when the continuum model may be a poor representation of the biology (Figure 7). Especially in biological contexts where cell numbers are large but local fluctuations can play an important role, we prefer to adopt the approach of starting with a biologically motivated discrete model and carefully derive the associated continuum limit, since this approach explicitly highlights the underlying assumptions inherent in the continuum model and provides us with a way of testing the accuracy of such assumptions.

## 4 Conclusion

In this work, we present a new one-dimensional cell-based model of heterogeneous epithelial tissue mechanics that includes cell proliferation and death. The main focus is to determine the corresponding continuum model which is a novel coupled system of nonlinear partial differential equations. The cell density equation is a parabolic partial differential equation while the cell property equations are hyperbolic partial differential equations. In deriving the continuum model, the discrete mechanisms and assumptions that underpin the continuum model have been made explicit by presenting the details of the coarse-graining derivation. Assumptions that relate to mean-field approximations and statistical independence of quantities are normally implicitly assumed in continuum models. By specifying the details of the derivation, and all assumptions required, our work provides insight into situations when these assumptions hold, as well as giving insight into when these assumptions fail to hold, such as when the number of cells, *N*(*t*), is not sufficiently large, when cell properties vary rapidly in space, when mechanical relaxation is slow relative to rate of proliferation, or with proliferation and death mechanisms which vary rapidly with respect to cell-length. Under these conditions we recommend that the discrete description is more appropriate than the approximate continuum description. Further, we stress the limitations of developing continuum models by simply adding source and sink terms to an existing model without considering the underlying discrete model in complex biological systems.

By coupling mechanics with proliferation and death we are able to explore biological scenarios that could not be described in previous modelling frameworks. Specifically we can focus on mechanical cell competition driven by variations in cell stiffness and resting cell length. By choosing mechanical relaxation rates sufficiently fast relative to proliferation rates we observe good agreement between the average of many identically prepared stochastic realisations of the discrete model and the corresponding solutions of the continuum model. This holds even when our simulations only consider 40 cells which is extremely small in comparison to the number of cells in an epithelial tissue. A continuum model is beneficial as we now have a tissue-level understanding of the mechanisms encoded in the discrete model and the time to solve the continuum model is independent of *N*(*t*). The discrete model remains beneficial and can provide additional information. For example, the average of many discrete realisations can match the continuum model but every discrete realization could go extinct which is not observed in the continuum model.

We explore mechanical cell competition applied to cancer invasion by considering cancer cells adjacent to healthy cells which compete for space. Interestingly, when we only allow cancer cells and healthy cells to differ in their cell stiffnesses, as a result of mechanical coupling, we observe that the cancer cells have more opportunities to proliferate and are less likely to die than healthy cells. We can then identify the cancer cells, as a result of the property of lower cell stiffness, as being the *winner* cells which invade the full domain. The influence of cell stiffness and cell size may therefore be an important factor to include when interpreting proliferation and death rates in experimental data. This analysis would not be possible using other existing models.

In all simulations we set *a* = 0 to model cells being in extension (Wyatt et al. 2016). Setting *a* > 0 gives qualitatively very similar results for homogeneous populations and also for heterogeneous populations when cells remain in extension throughout the simulation. This modelling framework is well-suited to be extended to cases where cells may also become compressed, for example in a tumour spheroid (Delarue et al. 2014). The model is well-suited to also study other observations of melanoma tumour spheroids such as subpopulations with differing proliferation rates located in different regions of the tumour, cells switching between these subpopulations, and the role of oxygen and nutrient concentrations (Haass 2015, Vittadello et al. 2020).

Many interesting extensions to this work are possible. Mathematically, the extent to which the continuum-limit holds with a free boundary is not yet clear. A free boundary also allows us to consider tissue growth (Serra-Picamal et al. 2012) and shrinkage in mechanically less constrained environments, such as in developmental biology. Further, explicitly incorporating additional biological mechanisms that regulate cell size (Holmes et al. 2016, Zmurchok et al. 2018, Zmurchok et al. 2020) and the evolution of intrinsic cell properties (Han et al. 2019) would be both mathematically interesting and biologically relevant. The theoretical foundations presented here for building a discrete model and constructing the continuum limit of that discrete model could be used to describe these additional mechanisms in future analyses.

This work was funded by the Australian Research Council (DP170100474). R.E.B is a Royal Society Wolfson Research Merit Award holder, would like to thank the Leverhulme Trust for a Research Fellowship and also acknowledges the BBSRC for funding via grant no. BB/R000816/1.

## Data Access

This article does not contain any additional data. Key algorithms used to generate results are available on https://github.com/ryanmurphy42/Murphy2020a.git.

## Author Contributions

All authors conceived and designed the study; R.J.M. performed numerical simulations and drafted the article; all authors provided comments and gave final approval for publication.

