## Supplementary Material for "Mechanical cell competition in heterogeneous epithelial tissues"

#### Contents

**Page No.**

SM1 Model formulation 3

SM1.1 Discrete model with $m > 1$ springs per cell 3

SM1.2 Death at boundaries 5

SM1.3 Derivation of proliferation with $m > 1$ springs per cell 6

SM1.4 Mechanical relaxation 11

SM2 Numerical methods 12

SM2.1 Discrete model 12

SM2.2 Continuum model 13

SM3 Homogeneous population 17

SM3.1 Extinction for constant proliferation and death 17

SM3.2 Reduced variance with cell-length-dependent mechanisms 19

SM3.3 Logistic proliferation and death 21

SM3.4 Piecewise proliferation: varying mechanical relaxation rate 23

SM4 Mechanical cell competition 26

SM4.1 Two populations: cell size at mechanical equilibrium 26

SM4.2 Logistic proliferation and death mechanisms 27

SM4.3 Varying mechanical relaxation rate 30

### S1 Model formulation

In Section 2.1 of the main manuscript, we present the discrete model with one spring per cell and the derivation of the corresponding continuum model. Here, we present the discrete model with $m > 1$ springs per cell, see Supplementary Material Section S1.1. This is important to define the field functions in the continuum model, in particular the mechanical relaxation term, for sufficiently small $N$ (Murphy et al. 2019) where the size of a cell is no longer small in comparison to the size of the domain. Proliferation still occurs at a cellular level rather than at a spring level with $m > 1$ therefore we still require many cells, i.e. $a_i \ll \delta x \ll L$. We also present: the special cases we need to consider at the boundaries for cell death, see Supplementary Section S1.2; the derivation of continuum model for proliferation with $m > 1$ springs per cell, see Supplementary Material Section S1.3; and in Supplementary Material Section S1.4 highlight key points for the derivation of the mechanical relaxation term (Murphy et al. 2019).

#### S1.1 Discrete model with $m > 1$ springs per cell

The model described in the main manuscript is presented by considering each cell to be represented by a single mechanical spring, $m = 1$, and tracking the evolution of cell boundaries. We now replace each cell with $m > 1$ identical springs (Figure S1a) (Murphy et al. 2019). We have spring boundaries at the cell boundaries as before but now we also have spring boundaries internal to the cell. We now track the evolution of all spring boundaries. In a system of $N$ cells, cell $i$ has spring boundaries $x_{i,\nu}^N, \nu = 1, \ldots, m$, where $x_i^N = x_{i,1}^N$. The spring length is defined as $l_{i,\nu}^N = x_{i,\nu}^N - x_{i,\nu-1}^N > 0$ and is related to cell length through $l_i^N \sim m l_{i,\nu}^N$ as $m \to \infty$, and with equality for all $m$ as $t \to \infty$. The viscosity coefficient for a cell, $\eta$, and mechanical cell properties, $k_i$ and $a_i$, are related to viscosity coefficient for a spring boundary, $\eta_\nu$, spring stiffness, $k_{i,\nu}^N$ and resting spring length, $a_{i,\nu}^N$, through the following scalings

$$\eta_\nu = \frac{\eta}{m}, \quad k_{i,\nu}^N = m k_i^N, \quad a_{i,\nu}^N = \frac{a_i^N}{m}, \tag{1}$$

The spring boundaries, $x_{i,\nu}^N$, evolve according to

$$\begin{aligned} \eta_\nu \frac{\mathrm{d}x_{i,\nu}^N(t)}{\mathrm{d}t} &= f_{i,\nu}^N - f_{i,\nu-1}^N, \quad i = 2, \ldots, N, \quad \nu = 1, 2, \ldots, m, \\ f_{i,\nu}^N &= k_{i,\nu}^N \left( l_{i,\nu}^N - a_{i,\nu}^N \right). \end{aligned} \tag{2}$$

We consider proliferation to be a property of a cell rather than a property of springs within a cell. Specifically, when spring $\nu$ in cell $i$ is chosen to proliferate we consider that the whole cell proliferates and introduce an additional $m$ springs (Figure S1b). Accordingly, we will introduce scaled spring proliferation rates. As with $m = 1$, we introduce the new cell boundary at the midpoint of original cell before proliferation. However, now we introduce $m$ additional springs. To do so we equally space $2m$ springs within the original cell. Similarly, for cell death we now instantly coalesce the cell boundaries and all internal spring boundaries to the centre of the dying cell. We have spring proliferation and death laws, $P_\nu$ and $D_\nu$, respectively,

$$P_\nu\left(l_{i,\nu}^N\right) = \frac{P\left(l_i^N\right)}{m}, \quad D_\nu\left(l_{i,\nu}^N\right) = \frac{D\left(l_i^N\right)}{m}. \tag{3}$$

The scalings are chosen such that the cell boundary velocities and proliferation/death rates are maintained and are independent of $m$ (Murphy et al. 2019).

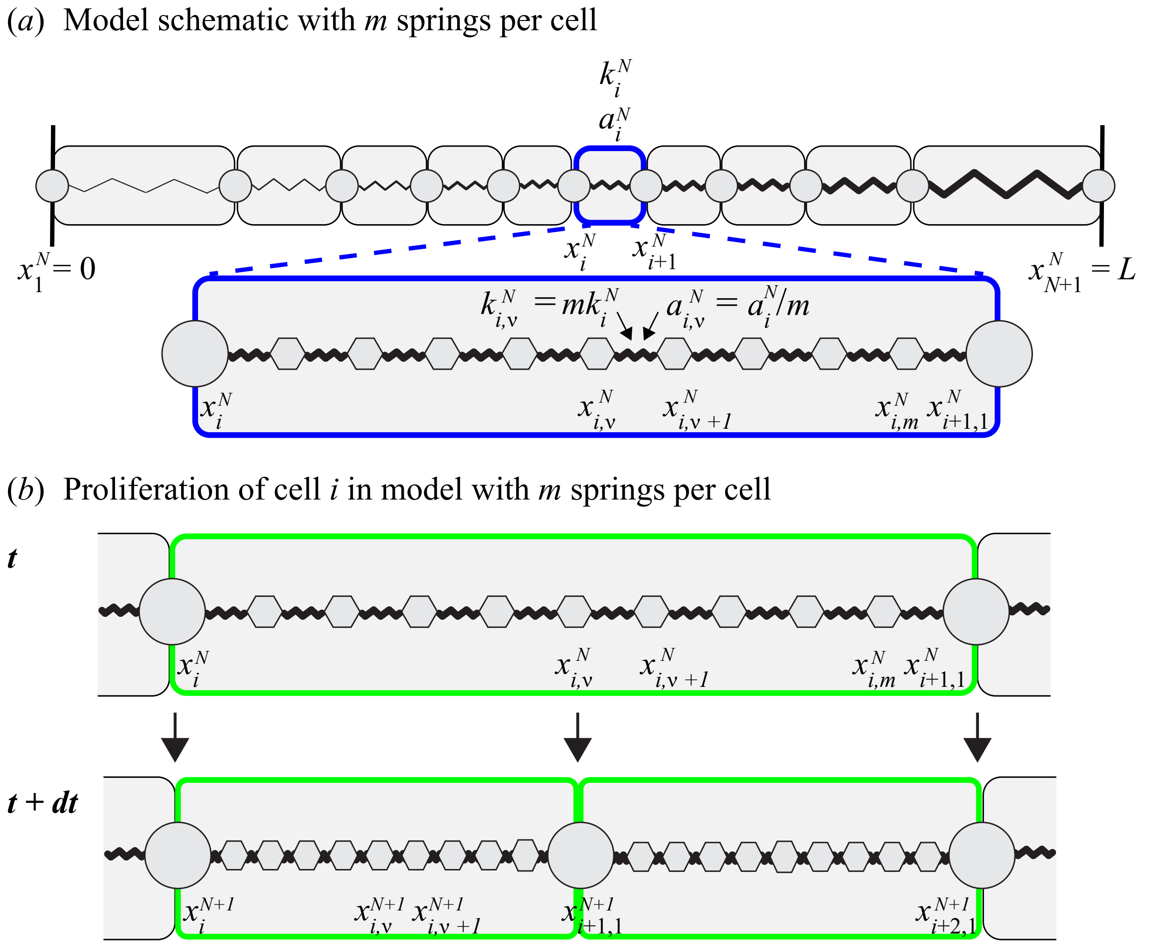


Figure S1: (a) Schematic for the discrete model with $m$ springs per cell. Heterogeneous population with $N$ cells in a fixed domain of length $L$. Cell $i$, with a blue border, is prescribed with cell stiffness $k_i^N$ and resting cell length $a_i^N$. The length of cell $i$ is $l_i^N = x_{i+1}^N - x_i^N$. Spring $\nu$ in cell $i$ has spring boundaries $x_{i,\nu}^N, x_{i,\nu+1}^N$. Each spring is prescribed with a spring stiffness $k_{i,\nu}^N = mk_i^N$ and a resting spring length $a_{i,\nu}^N = a_i^N/m$. Each spring has spring length $l_{i,\nu}^N = x_{i,\nu+1}^N - x_{i,\nu}^N$. The cell and spring boundaries are shown as discs and hexagons, respectively. (b) Proliferation of cell $i$, with a green border, in a model with $m$ springs per cell. The original cell divides into two cells. An additional $m$ equally spaced springs are introduced. This schematic is presented for even $m$.

#### S1.2 Death at boundaries

Cell death at a boundary is a special case of the discrete model which we present here. For death of the first cell whose left boundary is at $x = 0$, the first cell is removed and the left boundary of the second cell is set to $x = 0$ (Figure S2a). Similarly, for the death of the last cell whose right boundary is at $x = L$, the final cell is removed and the right boundary of cell $N - 1$ is set to $x = L$ (Figure S2b).

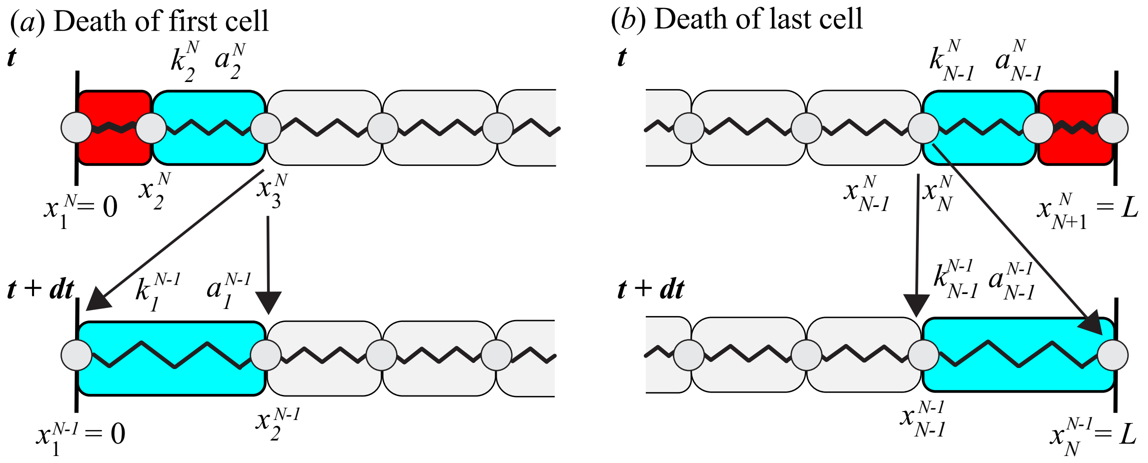


Figure S2: Model schematic for special cases of cell death of (a) first and (b) last cell.

#### S1.3 Derivation of proliferation with $m > 1$ springs per cell

In Section 2.2 of the main manuscript, we present the derivation with one spring per cell, $m = 1$. Here, for completeness, we rewrite the derivation with $m$ springs per cell. The advantage of $m$ springs per cell is that we can more appropriately define the continuous field functions, in particular the mechanical relaxation term for low cell numbers. New text relevant for $m$ springs per cell is shown in blue. Setting $m = 1$ in this new derivation recovers the results in the main manuscript.

Starting from the discrete model described in Section 2.1 but now for $m$ springs per cell as described in Supplementary Material Section 1.1, we now derive the proliferation source term in Equation (3). As cell proliferation is included stochastically, we consider an infinitesimal time interval $[t, t+\mathrm{d}t)$ and condition on the possible proliferation events that could occur and influence the position of spring boundary $\nu$ in cell $i$ in a system of $N$ cells. We note that proliferation is still considered a cell event rather than a spring event. Specifically, we say that a cell proliferates once any spring in the cell has been chosen to proliferate; this is captured through the scalings in Equation (3).

Choosing $\mathrm{d}t$ sufficiently small so that at most one proliferation event can occur in $[t, t+\mathrm{d}t)$ there are five possibilities: i) either there is no proliferation, in which case the spring boundary position ${x_{i,\nu}}^N$ only changes by mechanical relaxation; ii) there is proliferation to the right of cell $i$; iii) there is proliferation to the left of cell $i-1$; iv) cell $i-1$ proliferates; and v) cell $i$ proliferates. This leads to the following infinitesimal evolution law for the position of spring boundary $\nu$ in cell $i$ $x_{i,\nu}^N$, accounting for cell relabelling when a new cell is added:

$$
\begin{aligned}
{x_{i,\nu}}^N(t+\mathrm{d}t) = &\left[{x_{i,\nu}}^N(t) + \frac{\mathrm{d}t}{\eta}\left\{f^N\left({l_{i,\nu}}^N\right) - f^N\left({l_{i,\nu-1}}^N\right)\right\}\right] \times \mathbb{1}\left\{\text{no proliferation}\right\} \\
&+ \left[{x_{i,\nu}}^{N-1}(t) + \frac{\mathrm{d}t}{\eta}\left\{f^{N-1}\left({l_{i,\nu}}^{N-1}\right) - f^{N-1}\left({l_{i,\nu-1}}^{N-1}\right)\right\}\right] \\
&\qquad \times \mathbb{1}\left\{\text{proliferation right of cell } i\right\} \\
&+ \left[{x_{i-1,\nu}}^{N-1}(t) + \frac{\mathrm{d}t}{\eta}\left\{f^{N-1}\left({l_{i-1,\nu}}^{N-1}\right) - f^{N}\left({l_{i-1,\nu-1}}^{N-1}\right)\right\}\right] \\
&\qquad \times \mathbb{1}\left\{\text{proliferation left of cell } i-1\right\} \\
&+ \left[x^{N-1}_{i-1,\left(\frac{m}{2}+\frac{\nu+1}{2}\right)}(t) + \frac{\mathrm{d}t}{\eta}\left\{f^{N-1}\left(l^{N-1}_{i-1,\left(\frac{m}{2}+\frac{\nu+1}{2}\right)}\right) - f^{N}\left(l^{N-1}_{i-1,\left(\frac{m}{2}+\frac{\nu-1}{2}\right)}\right)\right\}\right] \\
&\qquad \times \mathbb{1}\left\{\text{proliferation of cell } i-1\right\}. \\
&+ \left[x^{N-1}_{i,\left(\frac{\nu+1}{2}\right)}(t) + \frac{\mathrm{d}t}{\eta}\left\{f^{N-1}\left(l^{N-1}_{i+1,\left(\frac{\nu+1}{2}\right)}\right) - f^{N}\left(l^{N-1}_{i-1,\left(\frac{\nu+1}{2}\right)}\right)\right\}\right] \\
&\qquad \times \mathbb{1}\left\{\text{proliferation of cell } i\right\}.
\end{aligned}
\tag{4}
$$

The derivation is shown for even $m$ and odd $\nu$ for simplicity (Figure S1b). With other choices of $m$ and $\nu$ we obtain slightly different terms for the proliferation of cell $i-1$ and $i$. However, these choices are not important to the following derivation. Each term in square brackets is the resulting force from neighbouring cells due to mechanical relaxation, given by Equations (2), for each potential event. In addition, we include Boolean variables expressed as indicator functions, $\mathbb{1}\{\cdot\}$, defined as

$$\mathbb{1}\{\text{event}\} = \begin{cases} 1, & \text{if event occurs in } [t, t+\mathrm{d}t), \\ 0, & \text{otherwise,} \end{cases} \tag{5}$$

whose expectations in the context of Equation (4) can be interpreted as proliferation probabilities. For a system of $N$ cells with $m$ springs per cell, where $\mathrm{d}t$ is sufficiently small, these proliferation probabilities are given by

$$\mathbb{P}\left(\text{no proliferation in } [t, t+\mathrm{d}t)\right) = 1 - \mathrm{d}t \sum_{j=1}^{N} \sum_{\nu=1}^{m} P_\nu\left(l_{j,\nu}{}^N\right), \tag{6a}$$

$$\mathbb{P}\left(\text{proliferation to the right of cell } i \text{ in } [t, t+\mathrm{d}t)\right) = \mathrm{d}t \sum_{j=i+1}^{N} \sum_{\nu=1}^{m} P_\nu\left(l_{j,\nu}{}^N\right), \tag{6b}$$

$$\mathbb{P}\left(\text{proliferation to the left of cell } i-1 \text{ in } [t, t+\mathrm{d}t)\right) = \mathrm{d}t \sum_{j=1}^{i-2} \sum_{\nu=1}^{m} P_\nu\left(l_{j,\nu}{}^N\right), \tag{6c}$$

$$\mathbb{P}\left(\text{proliferation of cell } i-1 \text{ in } [t, t+\mathrm{d}t)\right) = \mathrm{d}t \sum_{\nu=1}^{m} P_\nu\left(l_{i-1,\nu}{}^N\right), \tag{6d}$$

$$\mathbb{P}\left(\text{proliferation of cell } i \text{ in } [t, t+\mathrm{d}t)\right) = \mathrm{d}t \sum_{\nu=1}^{m} P_\nu\left(l_{i,\nu}{}^N\right), \tag{6e}$$

where using the proliferation rate scaling, from Equation (3), Equations (6) can be written in terms of the cell proliferation rates,

$$\mathbb{P}\left(\text{no proliferation in } [t, t+\mathrm{d}t)\right) = 1 - \mathrm{d}t \sum_{j=1}^{N} P\left(l_j^N\right), \tag{7a}$$

$$\mathbb{P}\left(\text{proliferation to the right of cell } i \text{ in } [t, t+\mathrm{d}t)\right) = \mathrm{d}t \sum_{j=i+1}^{N} P\left(l_j^N\right), \tag{7b}$$

$$\mathbb{P}\left(\text{proliferation to the left of cell } i-1 \text{ in } [t, t+\mathrm{d}t)\right) = \mathrm{d}t \sum_{j=1}^{i-2} P\left(l_j^N\right), \tag{7c}$$

$$\mathbb{P}\left(\text{proliferation of cell } i-1 \text{ in } [t, t+\mathrm{d}t)\right) = \mathrm{d}t\ P\left(l_{i-1}^N\right), \tag{7d}$$

$$\mathbb{P}\left(\text{proliferation of cell } i \text{ in } [t, t+\mathrm{d}t)\right) = \mathrm{d}t\ P\left(l_i^N\right). \tag{7e}$$

Taking a statistical expectation, denoted $\langle\cdot\rangle$, of Equation (4), $\langle x_{i,\nu}{}^N(t)\rangle$ now represents the expected position of spring boundary $\nu$ in cell $i$ at time $t$ in a system of $N$ cells. We use the proliferation probabilities with the following simplifying assumptions: i) $\left\langle x_{i,\nu}{}^N(t)\ \mathbb{1}\{\text{event}\}\right\rangle = \left\langle x_{i,\nu}{}^N(t)\right\rangle \langle \mathbb{1}\{\text{event}\}\rangle$, namely independence of the position of the cell boundary in space and proliferation propensity, and a mean-field approximation as proliferation propensities depend on spring length; ii) $\left\langle x_{i,\nu}{}^N(t)\ f^N(l_{j,\nu}{}^N)\right\rangle = \left\langle x_{i,\nu}{}^N(t)\right\rangle \left\langle f^N(l_{j,\nu}{}^N)\right\rangle$, namely independence of force and the propensity to proliferate, and a mean-field approximation as force depends on spring length; iii) a statistical mean-field approximation for force, $\langle f^N(l_{j,\nu}{}^N)\rangle = f^N\left(\langle l_{j,\nu}{}^N\rangle\right)$, and proliferation terms,

$\langle P(l_j^N)\rangle = P\left(\langle l_j^N\rangle\right)$. For simplicity, we now drop the $\langle\cdot\rangle$ notation. Then,

$$\begin{aligned}\frac{{x_{i,\nu}}^N(t+\mathrm{d}t)-{x_{i,\nu}}^N(t)}{\mathrm{d}t} &= \frac{1}{\eta}\left[f^N\left({l_{i,\nu}}^N\right)-f^N\left({l_{i,\nu-1}}^N\right)\right] \\ &- {x_{i,\nu}}^N(t)\sum_{j=1}^{N}P\left(l_j^N\right)+{x_{i,\nu}}^{N-1}(t)\sum_{j=i+1}^{N-1}P\left(l_j^{N-1}\right)+x^{N-1}_{i,\left(\frac{\nu+1}{2}\right)}(t)P\left(l_i^{N-1}\right) \\ &+ {x_{i-1,\nu}}^{N-1}(t)\sum_{j=1}^{i-2}P\left(l_j^{N-1}\right)+x^{N-1}_{i-1,\left(\frac{m}{2}+\frac{\nu+1}{2}\right)}(t)P\left(l_{i-1}^{N-1}\right)+\mathcal{O}\left(\mathrm{d}t\right).\end{aligned} \tag{8}$$

We also have

$$x^{N-1}_{i,\left(\frac{\nu+1}{2}\right)}(t)=x^{N-1}_{i,\nu}(t)+\left[x^{N-1}_{i,\left(\frac{\nu+1}{2}\right)}(t)-x^{N-1}_{i,\nu}(t)\right], \tag{9}$$

and we make assumption vii) that we are sufficiently far from the tissue boundary such that the term in square brackets of Equation (9) is negligible in comparison to the first term in the right hand side. This is consistent with Equation (16) in the derivation with $m=1$. Similarly for the term $x^{N-1}_{i-1,\left(\frac{m}{2}+\frac{\nu+1}{2}\right)}(t)$. This gives

$$\begin{aligned}\frac{{x_{i,\nu}}^N(t+\mathrm{d}t)-{x_{i,\nu}}^N(t)}{\mathrm{d}t} &= \frac{1}{\eta}\left[f^N\left({l_{i,\nu}}^N\right)-f^N\left({l_{i-1,\nu}}^N\right)\right] \\ &- {x_{i,\nu}}^N(t)\sum_{j=1}^{N}P\left(l_j^N\right)+{x_{i,\nu}}^{N-1}(t)\sum_{j=i+1}^{N-1}P\left(l_j^{N-1}\right)+x^{N-1}_{i,\nu}(t)P\left(l_i^{N-1}\right) \\ &+ {x_{i-1,\nu}}^{N-1}(t)\sum_{j=1}^{i-2}P\left(l_j^{N-1}\right)+x^{N-1}_{i-1,\nu}(t)P\left(l_{i-1}^{N-1}\right)+\mathcal{O}\left(\mathrm{d}t\right)\end{aligned} \tag{10}$$

We also assume: iv) the total propensity to proliferate is not significantly changed due to a single proliferation event, $\sum_{j=1}^{N-1}P\left(l_j^{N-1}\right)\mathrm{d}t=\sum_{j=1}^{N}P\left(l_j^{N}\right)\mathrm{d}t+\mathcal{O}(\mathrm{d}t^2,\frac{1}{N})$; v) a single proliferation event does not significantly alter the position of a cell boundary, ${x_{i,\nu}}^{N-1}(t)={x_{i,\nu}}^{N}(t)+\mathcal{O}(\mathrm{d}t)$. As we will show, assumptions iv) and v) are good approximations for large $N$ and allow us to combine summations. Then, assuming vi) $\langle {x_{i,\nu}}^N(t)\rangle$ is a continuous function of time, we rearrange and take the limit $dt\to 0$. For the proliferation terms we replace the cell length with the discrete cell density $q_i^N=1/l_i^N$ to obtain

$$\begin{aligned}\frac{\mathrm{d}{x_{i,\nu}}^N}{\mathrm{d}t} =& f^N\left({l_{i,\nu}}^N\right)-f^N\left({l_{i,\nu-1}}^N\right) \\ & -\left(\frac{1}{q_i^N(t)}\right)\left[\sum_{j=1}^{i-1}P\left(\frac{1}{q_{j+1}^N(t)}\right)\right].\end{aligned} \tag{11}$$

Equation (11) is only valid for the time interval $[t,t+\mathrm{d}t)$ under the assumptions iv) and v) above.

Thus far, we have extended the discrete model with mechanical relaxation to include the effects of cell proliferation and death. However, the statistically averaged model still retains information about discrete cell entities. We thus average over space to define a continuum cell density. Following Murphy et al. (Murphy et al. 2019), we introduce the microscopic density of cells,

$$\hat{q}(x,t)=\frac{1}{m}\sum_{i=1}^{N}\sum_{\nu=1}^{m}\delta\left(x-{x_{i,\nu}}^N(t)\right), \tag{12}$$

where $\delta$ is the Dirac delta function (Evans et al. 2008, Lighthill 1958). We define a local spatial average over a length scale $\delta x$, denoted $\langle\cdot\rangle_{\delta x}$, such that $a_{i,\nu} \ll a_i \ll \delta x \ll L$, which is sufficiently large to capture local heterogeneities for cellular properties that are constant during cell motion, including $k$ and $a$, but sufficiently small to define continuous properties across $L$. The continuous cell density function, $q(x,t)$, is thus defined as

$$q(x,t) = \langle \hat{q}(x,t) \rangle_{\delta x} = \frac{1}{2\delta x} \int_{x-\delta x}^{x+\delta x} \hat{q}(y,t)\ \mathrm{d}y. \tag{13}$$

Differentiating Equation (13) with respect to time gives

$$\frac{\partial q(x,t)}{\partial t} = -\frac{\partial}{\partial x} \left\langle \frac{1}{m} \sum_{i=1}^{N} \sum_{\nu=1}^{m} \delta\left(x - {x_{i,\nu}}^{N}(t)\right) \frac{\mathrm{d}{x_{i,\nu}}^{N}}{\mathrm{d}t} \right\rangle_{\delta x}, \tag{14}$$

where we use properties of the Dirac delta distribution (Lighthill 1958) and interchange the derivative with the spatial average as $\delta x$ is small. Consistent with assumptions 4)-5) above, the sum over the microscopic densities can be considered to be fixed over $N$ cells in Equation (14) within the small time interval $[t, t + \mathrm{d}t)$.

On the right hand side of Equation (11), the first two terms involving $f$ correspond to a mechanical contribution. This contribution is unchanged compared to Murphy et al. (Murphy et al. 2019) and, when substituted into Equation (14), it gives rise to the mechanical relaxation term on the right hand side of the continuum equation (3) (Supplementary Material S1.4). We now focus only on the contribution determined by substituting the proliferation terms of Equation (11) into Equation (14), giving a contribution which we denote $\partial q(x,t)/\partial t|_P$,

$$\left.\frac{\partial q(x,t)}{\partial t}\right|_P = \frac{\partial}{\partial x} \left\langle \frac{1}{m} \sum_{i=1}^{N} \sum_{\nu=1}^{m} \delta\left(x - {x_{i,\nu}}^{N}(t)\right) \frac{1}{q_{i-1}^{N}(t)} \left[ \sum_{j=1}^{i-1} P\left(\frac{1}{q_j^N(t)}\right)\right] \right\rangle_{\delta x}. \tag{15}$$

At this point in the derivation for $m = 1$ we make the assumption that we are sufficiently far from the tissue boundary. We name this as assumption vii). For $m > 1$ we have already made assumption vii) and this is not required again here. To switch the dependence on the cell index to cell position, we focus on the sum in square brackets on the right hand side of Equation (15). We multiply each term $j$ by $1 = l_j q_j$. Then relating the discrete cell density to the continuous density through $q_j^N = q(x_j^N(t), t)$ gives

$$\sum_{j=1}^{i-1} q(x_j^N(t), t) P\left(\frac{1}{q(x_j^N(t), t)}\right) l_j. \tag{16}$$

We discretise the spatial domain $x_1 \leq x \leq x_{i-1}$ with a uniform mesh with nodes $y_s, s = 1, 2, \ldots, S$, where $y_1 = x_1$, $y_S = x_{i-1}$, and $y_s - y_{s-1} = \Delta y \ll l_j$. Then, evaluating the continuous density at each node position, $y_s$, we interpret Equation (16) as the following Riemann sum

$$\sum_{s=1}^{S} q(y_s, t) P\left(\frac{1}{q(y_s, t)}\right) \Delta y = \int_0^{x_i^N} q\left(y, t\right) P\left(\frac{1}{q(y,t)}\right)\ \mathrm{d}y. \tag{17}$$

where the integral on the right hand side is obtained by taking the limit $\Delta y \to 0$. Substituting Equation (17) into Equation (15) gives

$$\left.\frac{\partial q(x,t)}{\partial t}\right|_P = \frac{\partial}{\partial x}\left\langle \frac{1}{m}\sum_{i=1}^{N}\sum_{\nu=1}^{m}\delta\left(x - x_{i,\nu}{}^{N}(t)\right)\left(\frac{1}{q_{i-1}^N}\right)\left[\int_0^{x_i^N} q\left(y,t\right)P\left(\frac{1}{q(y,t)}\right)\,\mathrm{d}y\right]\right\rangle_{\delta x}. \quad (18)$$

Calculating the spatial average, which only includes contributions from within the spatial average interval due to the Dirac delta functions, gives

$$\left.\frac{\partial q(x,t)}{\partial t}\right|_P = \frac{\partial}{\partial x}\left(\left(\frac{n}{2\delta x}\right)\frac{1}{n}\frac{1}{m}\sum_{r=1}^{n}\sum_{\nu=1}^{m}\left(\frac{1}{q_{r-1}^N}\right)\left[\int_0^{x_r^N} q\left(y,t\right)P\left(\frac{1}{q(y,t)}\right)\,\mathrm{d}y\right]\right), \quad (19)$$

where the index $r$ labels the $n$ cell boundaries contained within the spatial average interval $(x - \delta x, x + \delta x)$. Equation (19) is now independent of $m$, which is to be expected as proliferation is considered a cell event rather than a spring event. Simplifying gives

$$\left.\frac{\partial q(x,t)}{\partial t}\right|_P = \frac{\partial}{\partial x}\left(\left(\frac{n}{2\delta x}\right)\frac{1}{n}\sum_{r=1}^{n}\left(\frac{1}{q_{r-1}^N}\right)\left[\int_0^{x_r^N} q\left(y,t\right)P\left(\frac{1}{q(y,t)}\right)\,\mathrm{d}y\right]\right). \quad (20)$$

Since $a_i \ll \delta x \ll L$ and $n \gg 1$ we have $q_r^N = q(x_r^N(t),t) = q(x,t)$ for all $r$, which is independent of $r$. Similarly, $x_r \approx x$ for all $r$, where $x$ is the centre of the spatial average interval. This gives

$$\left.\frac{\partial q(x,t)}{\partial t}\right|_P = \frac{\partial}{\partial x}\left(\left(\frac{n}{2\delta x}\right)\frac{1}{q(x,t)}\int_0^{x} q\left(y,t\right)P\left(\frac{1}{q(y,t)}\right)\,\mathrm{d}y\right). \quad (21)$$

As $n/(2\delta x) = q(x,t)$ in this spatial average interval, Equation (21) simplifies to

$$\left.\frac{\partial q(x,t)}{\partial t}\right|_P = q\left(x,t\right)P\left(\frac{1}{q(x,t)}\right). \quad (22)$$

At this point, we see that all explicit references to the total number of cells, $N(t)$, vanish. This allows the validity of the derivation, initially restricted to the time interval $[t, t+\mathrm{d}t)$, to be extended to arbitrary times. As $N(t) = \int_0^L q(x,t)\,\mathrm{d}x$, the change in the total cell number with time due to proliferation is accounted for through the source term written in Equation (22). We also stated assumption vii) that held true when sufficiently far from the tissue boundary but we find that this works at the boundary also (Section 2).

#### S1.4 Mechanical relaxation

We outline the key steps to derive the mechanical terms, with one spring per cell, and refer the reader to Section 2 of Murphy et al. (Murphy et al. 2019) for full details. We introduce field functions for the force, $f(x,t)$, cell stiffness, $k(x,t)$, and resting cell length $a(x,t)$, which relate to the discrete model through

$$\begin{aligned} f(x_i^N(t),t) &= f_i^N, \\ k(x_i^N(t),t) &= k_i^N, \\ a(x_i^N(t),t) &= a_i^N. \end{aligned} \tag{23}$$

Substituting Equation (11) into Equation (14), we focus on the force term on the right side and consider no proliferation or death. We expand the cell-cell interaction force using the small cell length parameter $l_i^N$, small as the number of cell boundaries inside the spatial average interval is large, i.e. $n \gg 1$ in $(x - \delta x, x + \delta x)$. We then simplify to leading order, integrate over the spatial average interval, and perform spatial mean-field approximations using $n \gg 1$ in the spatial average interval. We arrive at the force term, $-(1/\eta)\, \partial^2 f/\partial x^2$, written in Equation (3), where the continuous cell-cell interaction force, $f$, is given by Equation (4). This force term has an important physical interpretation where the cell density flux, $j(x,t)$, is equal to the gradient of the cell-cell interaction force,

$$j(x,t) = \frac{1}{\eta}\frac{\partial f(x,t)}{\partial x}. \tag{24}$$

Further we find the cell velocity, $u(x,t)$, is related to the cell density and gradient of the cell-cell interaction force through Equation (5).

### S2 Numerical methods

Here we present the numerical methods used to solve the discrete and continuum models. Key algorithms used to generate results are available on GitHub (https://github.com/ryanmurphy42/Murphy2020a.git).

#### S2.1 Discrete model

We numerically solve the discrete model with a constant time step algorithm, using a forward Euler approximation to integrate the discrete equations (1), and rejection sampling to determine when proliferation and death events occur (Gelman et al. 2013). This method is valid for all proliferation and death law mechanisms which we consider. However, we note that to improve computational efficiency, in the case of the constant proliferation and death law mechanism, we could use Gillespie's algorithm (Gillespie 1977). This is possible as the propensities of cells to proliferate or die are constant within the calculated time to the next reaction interval. This is more difficult for the linear and logistic proliferation and death mechanisms where, due to mechanical coupling, the propensity of a cell to proliferate or die can vary appreciably within the calculated time to the next reaction interval per the Gillespie method. In such a case the Extrande method may be considered (Voliotis et al. 2016).

##### S2.1.1 Euler's method

To simulate a single discrete realization, we initialise the model with $N$ cells. We prescribe each cell $i$ with the mechanical cell properties including cell stiffness $k_i^N$ and resting cell length $a_i^N$. We prescribe proliferation and death mechanisms to each cell $i$ and any associated proliferation or death cell properties. We define the initial cell positions and then for each time step of size $\Delta t = 0.0001$ we update the cell positions using a simple forward Euler method to integrate Equations (1) numerically. At the end of each time step we determine whether a proliferation or death event occurs and if so which cell has proliferated or died. To do so we use rejection sampling (Gelman et al. 2013) where we generate three independent random numbers from a uniform distribution, $r_1, r_2, r_3 \sim U[0, 1]$. Then a cell event, which could be either a cell proliferation or cell death event, occurs when

$$r_1 < \sum_{i=1}^{N(t)} P(l_i^N)\Delta t + \sum_{i=1}^{N(t)} D(l_i^N)\Delta t,$$

i.e. with probability $\sum_{i=1}^{N(t)} \left[P(l_i^N) + D(l_i^N)\right] \Delta t$. Given that a cell event occurs, a proliferation event occurs if

$$r_2 < \frac{\sum_{i=1}^{N(t)} P(l_i^N)}{\sum_{i=1}^{N(t)} P(l_i^N) + \sum_{i=1}^{N(t)} D(l_i^N)}.$$

Otherwise we have a cell death event. To determine which cell is proliferating, similarly for dying, we find the index $j$ which satisfies,

$$\frac{\sum_{i=1}^{j} P(l_i^N)}{\sum_{i=1}^{N(t)} P(l_i^N)} < r_3 \leq \frac{\sum_{i=1}^{j+1} P(l_i^N)}{\sum_{i=1}^{N(t)} P(l_i^N)}. \tag{25}$$

We then update the node positions, cell properties, and indices according to the model description in Section 2.1. We repeat for each time step until we have reached the final time. This approach requires that at most one cell event can occur within each time step which is satisfied for the parameters used in this work. For other parameters where this assumption may not be valid the size of the time step could be reduced.

#### S2.2 Continuum model

We now outline the numerical method we use to solve the continuum model. First, for completeness, we rewrite the governing equations for cell density, $q(x,t)$,

$$\frac{\partial q(x,t)}{\partial t} = -\frac{1}{\eta}\frac{\partial^2 f(x,t)}{\partial x^2} + q(x,t)\ P\left(\frac{1}{q(x,t)}\right) - q(x,t)\ D\left(\frac{1}{q(x,t)}\right), \tag{26}$$

where the cell-cell interaction force, $f(x,t)$, is given by

$$f(x,t) = k(x,t)\left(\frac{1}{q(x,t)} - a(x,t)\right), \tag{27}$$

and $P(1/q(x,t)), D(1/q(x,t))$ are the proliferation and death mechanisms, respectively. The cell properties are governed by

$$\frac{\partial \chi(x,t)}{\partial t} + u(x,t)\frac{\partial \chi(x,t)}{\partial x} = 0, \quad \chi = k, a, \beta, \gamma, l_d, \tag{28}$$

where the cell velocity $u(x,t)$ is given by,

$$u(x,t) = \frac{1}{\eta q(x,t)}\frac{\partial f(x,t)}{\partial x}. \tag{29}$$

In Equation (26) for the cell density we have a second-order spatial derivative, and in the cell property equations, Equations (28), we have first-order spatial derivatives. Both equations have first-order time derivatives. To begin we discretise the domain of fixed length $L$ with a uniform mesh with spatial step $\Delta x$. We discretise time with a uniform mesh with time step $\Delta t$. Second-order spatial derivative terms are approximated by standard central differences. First-order spatial derivatives are approximated by standard upwind differences. Temporal derivatives are approximated by a Crank-Nicolson approximation. We then have a system of nonlinear algebraic equations for the cell density, cell stiffness, resting cell length, and any other cell properties specified by the proliferation and/or death laws. Each resulting system of nonlinear algebraic equations we solve sequentially within the same Newton-Raphson iteration (Chapra et al. 2010) until a convergence tolerance, $\epsilon$, is satisfied. In each iteration each resulting system of linearised tridiagonal algebraic equations is solved using the Thomas algorithm (Zheng et al. 2002). There are three key choices with this method $\Delta x$, $\Delta t$, and $\epsilon$. Any implementation of the numerical method should ensure that $\Delta x$, $\Delta t$, and $\epsilon$ are sufficiently small that the solution is grid-independent. In the results we present we take $\Delta x = 0.01$, $\Delta t = 0.00001$, and $\epsilon = 0.001$.

For convenience, we now explain the Newton-Raphson method in more detail and explicitly present how we discretise the cell density equation, its boundary condition, discretise cell property equations, their boundary conditions, and update the interface position when there are two adjacent populations.

##### S2.2.1 Newton-Raphson method

The following notation is used. We use the subscript $j = 1, 2, \ldots, J$ to represent spatial nodes. We use the superscript $n = 1, 2, \ldots, T$ to represent temporal nodes. We use the superscript $r = 0, 1, \ldots, R_n$ to represent the Newton-Raphson iterate within time step $n$, where iterate $r = R_n$ is the final iterate which meets the convergence tolerance $\epsilon$. For convenience, we will drop the $R_n$ notation for the final iterate of time step $n$, for example, we write $q_j^n = q_j^{n,R_n}$ for the cell density at spatial node $j$.

We now solve for each variable at time step $n+1$. The initial iterate, corresponding to $r = 0$, is given by $q_j^{n+1,0} = q_j^n$ and $\chi_j^{n+1,0} = \chi_j^n$ for $\chi = k, a, \beta, \gamma, l_d$. The following is for the $r^{\text{th}}$ iterate.

We first solve for the **cell density**. We substitute Equation (27) into Equation (26). For internal spatial nodes, $j = 2, 3, \ldots, J-1$, we have, after rearranging so that all terms are on the right hand side, and for convenience multiplying all terms by $\Delta t$, a system of algebraic equations

$$\begin{aligned}
0 = &- q_j^{n+1,r} + q_j^n \\
&- \frac{\Delta t}{2\eta(\Delta x)^2}\left[k_{j-1}^n\left(\frac{1}{q_{j-1}^n} - a_{j-1}^n\right) - 2k_j^n\left(\frac{1}{q_j^n} - a_j^n\right) + k_{j+1}^n\left(\frac{1}{q_{j+1}^n} - a_{j+1}^n\right)\right] \\
&- \frac{\Delta t}{2\eta(\Delta x)^2}\left[k_{j-1}^{n+1,r-1}\left(\frac{1}{q_{j-1}^{n+1,r}} - a_{j-1}^{n+1,r-1}\right) - 2k_j^{n+1,r-1}\left(\frac{1}{q_j^{n+1,r}} - a_j^{n+1,r-1}\right)\right. \\
&\qquad \left. + k_{j+1}^{n+1,r-1}\left(\frac{1}{q_{j+1}^{n+1,r}} - a_{j+1}^{n+1,r-1}\right)\right] \\
&+ \frac{\Delta t}{2}\left[q_j^n P\left(\frac{1}{q_j^n}\right) - q_j^n D\left(\frac{1}{q_j^n}\right)\right] \\
&+ \frac{\Delta t}{2}\left[q_j^{n+1,r} P\left(\frac{1}{q_j^{n+1,r}}\right) - q_j^{n+1,r} D\left(\frac{1}{q_j^{n+1,r}}\right)\right].
\end{aligned} \tag{30}$$

For boundary nodes, $j = 1, J$, we have fixed boundaries which correspond to zero velocity boundary conditions

$$\frac{\partial f(x,t)}{\partial x} = 0, \quad x = 0, L. \tag{31}$$

We apply a forward difference approximation to Equation (31) at $x = 0$, corresponding to spatial node $j = 1$,

$$0 = \frac{1}{\Delta x} \left[ k_2^{n+1,r-1} \left( \frac{1}{q_2^{n+1,r}} - a_2^{n+1,r-1} \right) - k_1^{n+1,r-1} \left( \frac{1}{q_1^{n+1,r}} - a_1^{n+1,r-1} \right) \right], \tag{32}$$

and a backward difference approximation to Equation (31) at $x = L$, corresponding to spatial node $j = J$,

$$0 = \frac{1}{\Delta x} \left[ k_J^{n+1,r-1} \left( \frac{1}{q_J^{n+1,r}} - a_J^{n+1,r-1} \right) - k_{J-1}^{n+1,r-1} \left( \frac{1}{q_{J-1}^{n+1,r}} - a_{J-1}^{n+1,r-1} \right) \right]. \tag{33}$$

Equations (30), (32) and (33) can be combined to form a tridiagonal matrix $\boldsymbol{F}(\boldsymbol{q^{n+1,r}})$, where $\boldsymbol{q}^{n+1,r}$ is the system of algebraic equations of the $r^{\text{th}}$ iterate of the Newton-Raphson method at time step $n+1$. We then calculate the Jacobian, $\boldsymbol{J}(\boldsymbol{q^{n+1,r}})$, of $\boldsymbol{F}(\boldsymbol{q^{n+1,r}})$. We form a linear system for the $r^{\text{th}}$ iterate of the Newton-Raphson method within the time step $n+1$ as

$$\boldsymbol{J}(\boldsymbol{q}^{n+1,r})\ \boldsymbol{\delta q}^{n+1,r} = -\boldsymbol{F}(\boldsymbol{q}^{n+1,r}), \tag{34}$$

where $\boldsymbol{\delta q}^{n+1,r}$ is the Newton-Raphson correction. As $\boldsymbol{F}(\boldsymbol{q}^{n+1,r})$ is tridiagonal we solve this using the Thomas algorithm (Zheng et al. 2002), to the determine the next iterate,

$$\boldsymbol{q}^{n+1,r+1} = \boldsymbol{q}^{n+1,r} + \boldsymbol{\delta q}^{n+1,r} \tag{35}$$

Next we sequentially solve for each **cell property**. We calculate the velocity at each node, $v_j^{n+1,r}$,

$$v_j^{n+1,r} = \frac{1}{\eta} \frac{1}{q_j^{n+1,r}} \left[ k_{j+1}^{n+1,r-1} \left( \frac{1}{q_{j+1}^{n,r}} - a_{j+1}^{n+1,r-1} \right) - k_{j-1}^{n+1,r-1} \left( \frac{1}{q_{j-1}^{n+1,r}} - a_{j-1}^{n+1,r-1} \right) \right]. \tag{36}$$

Then we substitute Equation (29) into Equation (28) and consider $\chi = k$. If $v_j^{n+1,r} > 0$, then we apply a backward difference approximation to the first-order spatial derivatives and a Crank-Nicolson approximation for the time derivative

$$\begin{aligned} 0 =& k_j^{n+1,r} - k_j^n \\ &- \frac{\Delta t}{2\eta(\Delta x)^2} \Bigg\{ \left[ k_j^n \left( \frac{1}{q_j^n} - a_j^n \right) - k_{j-1}^n \left( \frac{1}{q_{j-1}^n} - a_{j-1}^n \right) \right] \left[ k_j^n - k_{j-1}^n \right] \\ &+ \left[ k_j^{n+1,r} \left( \frac{1}{q_j^{n+1,r}} - a_j^{n+1,r-1} \right) - k_{j-1}^{n+1,r} \left( \frac{1}{q_{j-1}^{n+1,r}} - a_{j-1}^{n+1,r-1} \right) \right] \left[ k_j^{n+1,r} - k_{j-1}^{n+1,r} \right] \Bigg\}. \end{aligned} \tag{37}$$

Similarly, if $v_j^{n+1,r} < 0$, then we apply a forward difference approximation.

For only mechanical relaxation or for mechanical relaxation with proliferation, we have fixed boundary conditions at $x = 0, L$, at spatial nodes $j = 1, J$, respectively,

$$\begin{aligned} k_1^{n+1,r} &= k_1^n, \\ k_J^{n+1,r} &= k_J^n. \end{aligned} \tag{38}$$

These boundary conditions also apply for cell death in a homogeneous population. However, when we have cell death with two adjacent populations, for example in Section 3.2, we need to modify a boundary condition when one population becomes extinct. To do so, we calculate the total cell number for each tissue within each time step. When the total number of cells in a tissue decreases below one we remove this tissue by setting the interface position equal to the relevant domain boundary, and make cell properties homogeneous across the domain by setting them equal to the values in the remaining tissue. Then, similarly to Equations (34) and (35), we solve for $\boldsymbol{k}^{n+1,r}$.

In the examples presented in this work, the resting cell length, $a$, and proliferation and death properties, $\beta, \gamma$ and $l_d$, are homogeneous across the population. Therefore, they do not need to be simulated and we set $\boldsymbol{\delta a}^{n+1,r} = \boldsymbol{\delta \beta}^{n+1,r} = \boldsymbol{\delta \gamma}^{n+1,r} = \boldsymbol{\delta l_d}^{n+1,r} = 0$. However, if any of $a, \beta, \gamma$ or $l_d$ are heterogeneous, we extend the above by discretising the relevant cell property equations and boundary conditions similarly to Equations (37) and (38).

When we have two populations we also update the interface position, $s(t)$. To do so, we find the closest spatial node to $s(t)$ and calculate the velocity at this node. Suppose that the closest spatial node is at node $j$ then we set $\delta s^{n+1,r} = v_j^{n+1,r}\Delta t$ and update the interface position through

$$s^{n+1,r} = s^n + \delta s^{n+1,r}. \tag{39}$$

We iterate until $||\boldsymbol{\delta q}^{n+1,r}, \boldsymbol{\delta k}^{n+1,r}, \boldsymbol{\delta a}^{n+1,r}, \boldsymbol{\delta \beta}^{n+1,r}, \boldsymbol{\delta \gamma}^{n+1,r}, \boldsymbol{\delta l_d}^{n+1,r}, \delta s^{n+1,r}||_\infty < \epsilon$.

##### S2.2.2 Initial conditions

Equations (3)-(6) have the following initial conditions, for $0 < x < L$,

$$q(x,0) = q_0(x), \quad \chi(x,0) = \chi_0(x) \text{ for } \chi = k, a, \beta, \gamma, l_d. \tag{40}$$

If initial conditions are provided only for the discrete model they can be converted to continuum model initial conditions. This has been discussed in previous work, see Supplementary Section 1 of Murphy et al. 2019.

### S3 Homogeneous population

We now present additional results for the homogeneous population: an exact calculation for extinction with constant proliferation and death mechanisms, and results for logistic proliferation and death mechanisms.

#### S3.1 Extinction for constant proliferation and death

In Section 3.1 of the main manuscript, we consider a homogeneous population with constant proliferation and constant death mechanisms with equal rates. In this case, the average of the discrete realizations shows good agreement with the corresponding solution of the continuum model. However, each individual realization exhibits very different behaviour to the continuum model. Due to the total cell number following a linear birth-death process independent of mechanical relaxation, each individual realization will eventually become extinct and the averaged total cell number displays increasing variance with time (Ross 1996). We now present the exact expressions for the extinction probability, mean, and standard deviation of the linear birth-death process. Corresponding results from the discrete realizations are then shown to match these expressions.

For the constant proliferation and death law combination with equal proliferation and death rates, $\beta$, we follow the work of Morgan (1977) who applies conditioning arguments to find the following probability generating function

$$G(z;t) = \left(\frac{z + \beta t\,(1-z)}{1 + \beta t\,(1-z)}\right)^{N(0)}, \tag{41}$$

where $N(0)$ is the initial cell population and $z$ is a dummy variable defined for $|z| \leq 1$. The extinction probability, $\mathbb{P}\,(E)$, is

$$\mathbb{P}\,(E) = \mathbb{P}\,(N(t)=0) = G(0;t) = \left(\frac{\beta t}{1+\beta t}\right)^{N(0)}. \tag{42}$$

We observe that $\mathbb{P}\,(N(t)=0) \to 1$ as $t \to \infty$, i.e. every individual realization will eventually become extinct. The mean is the same as the initial cell number,

$$\mathbb{E}\,(N(t)) = \left.\frac{\partial G(z;t)}{\partial z}\right|_{z=1} = N(0). \tag{43}$$

The standard deviation, $\sigma(t)$, is

$$\sigma(t) = \sqrt{2N(0)\beta t}. \tag{44}$$

We now compare these exact formulas with our discrete simulation, with $N(0) = 40$ and find very good agreement (Figure S3.1).

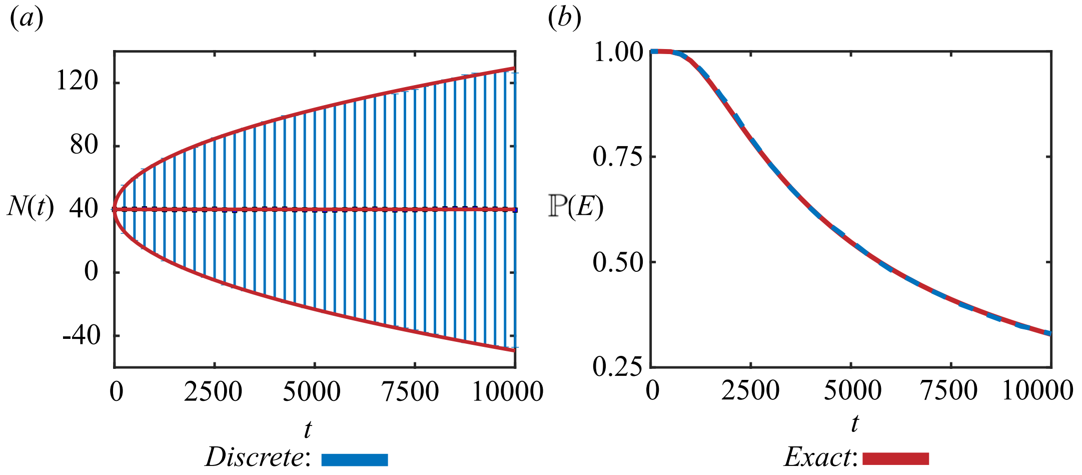


Figure S3: Comparison of 5000 discrete realizations with the exact expressions for the (a) mean and variance of total cell number and the (b) probability of extinction, $\mathbb{P}(E)$. Results shown for a homogeneous population with $k = 10$, $a = 0$, constant proliferation and death laws with equal rates, $\beta = 0.01$, and $0 < t < 10000$.

##### S3.2 Reduced variance with cell-length-dependent mechanisms

In Figures 3 and 4 in the main manuscript, we observe reduced variance in population with cell-length dependent proliferation and death mechanisms in comparison to cell-length independent proliferation and death mechanisms. Here, we explore the difference between constant and linear proliferation death mechanisms by presenting a single realisation for each (Figure S4). In the constant proliferation and death mechanism case the net proliferation rate, which is the sum of probabilities of each cell to proliferate minus the sum of probabilities of each cell to die, is always zero (Figure S4e). However, with the linear proliferation and death mechanism the net proliferation rate adjusts, due to changes in number of cells and their cell lengths, to stabilise the population at its equilibrium value (Figure S4f,h).

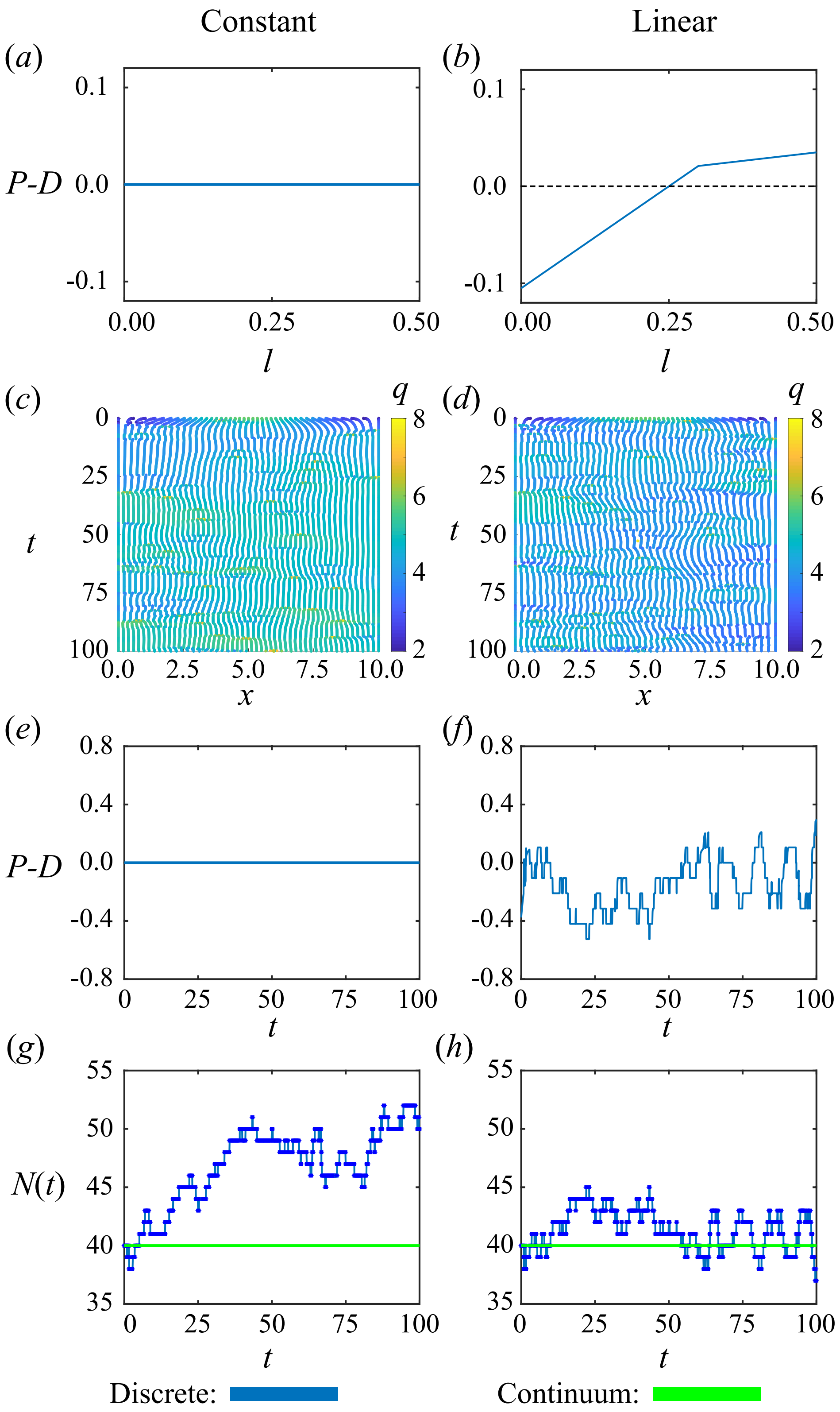


Figure S4: Homogeneous population with constant and linear cell-length dependent proliferation and death mechanisms. (a)-(b) Net proliferation rate for a single cell, $P - D$, dependent cell length, $l$. (c)-(d) Single realizations of cell boundary characteristics for $0 \leq t \leq 100$. (e)-(f) Net proliferation rate for the single realizations. (g)-(h) Total cell number for single realization (blue) compared to the continuum solution (green).

#### S3.3 Homogeneous population: logistic proliferation and death

In Figures 3 and 4 in the main manuscript we present results for a homogeneous population with constant and linear cell-length-dependent proliferation mechanisms, respectively. We now present results for homogeneous population for the logistic proliferation and death mechanism (Figure S5).

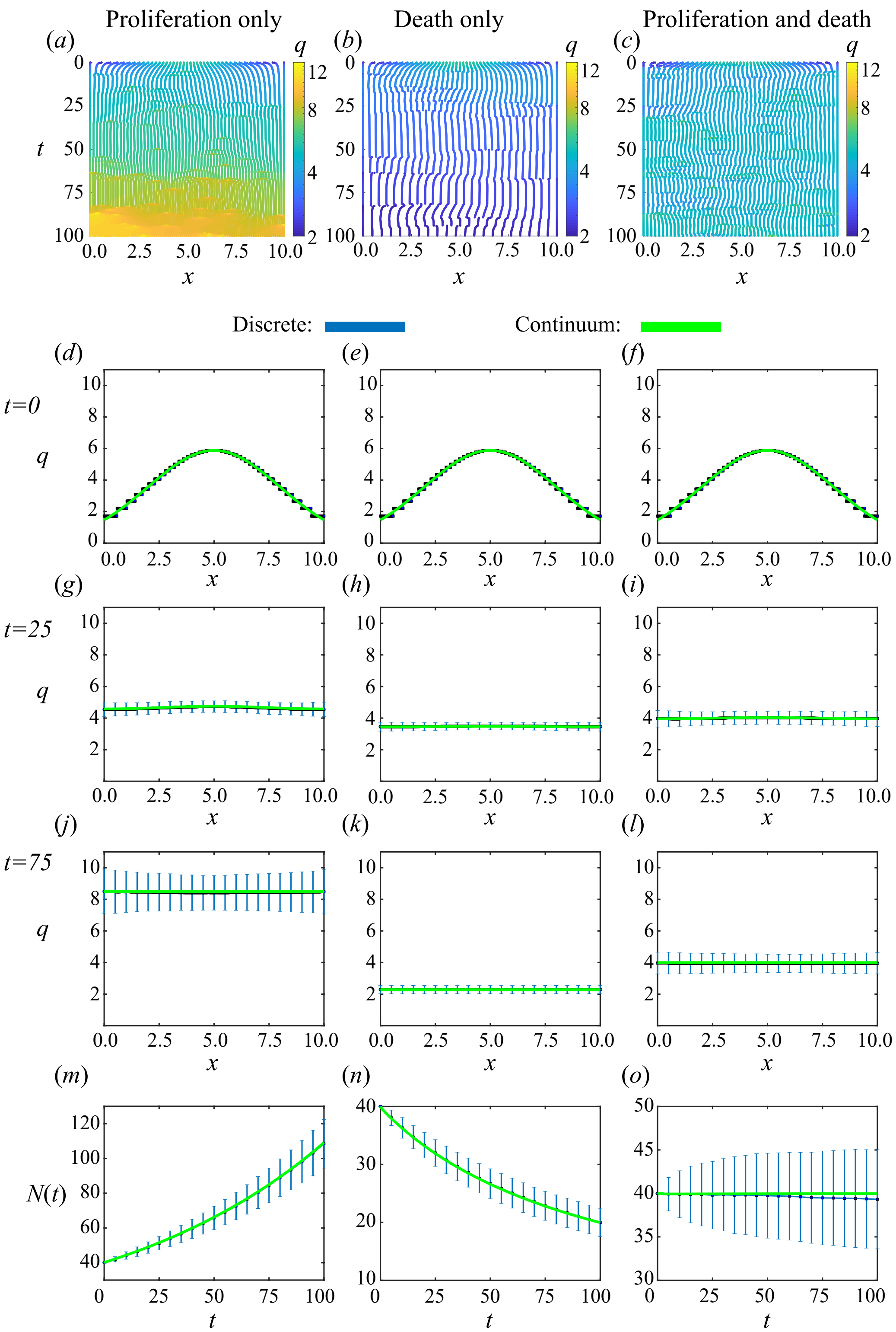


Figure S5: Homogeneous population with **logistic** proliferation and death mechanisms. Proliferation only, death only, and proliferation with death shown in left, middle and right columns, respectively. (a)-(c) Single realizations of cell boundary characteristics for $0 \leq t \leq 100$. (d)-(f), (g)-(i), (j)-(l) Density snapshots at times $t = 0, 25, 75$, respectively. (m)-(o) Total cell number. The average and standard deviation (blue error bars) of 2000 discrete simulations are compared to solution of continuum model (green).

##### S3.4 Piecewise proliferation: varying mechanical relaxation rate

In Figure 7 in the main manuscript we show that the solution of the continuum model can differ significantly from the solution to the discrete model with slow mechanical relaxation, $k = 0.0001$, and matches extremely well when $k = 1000$. We now present the density snapshots corresponding to Figure 7. For $k = 0.0001$ the density snapshots have jumps at locations of the initial cell boundaries and density in the discrete model is higher than the continuum model (Figure S6a,c,e). For $k = 1000$ the discrete model and continuum model are consistent (Figure S6b,d,f).

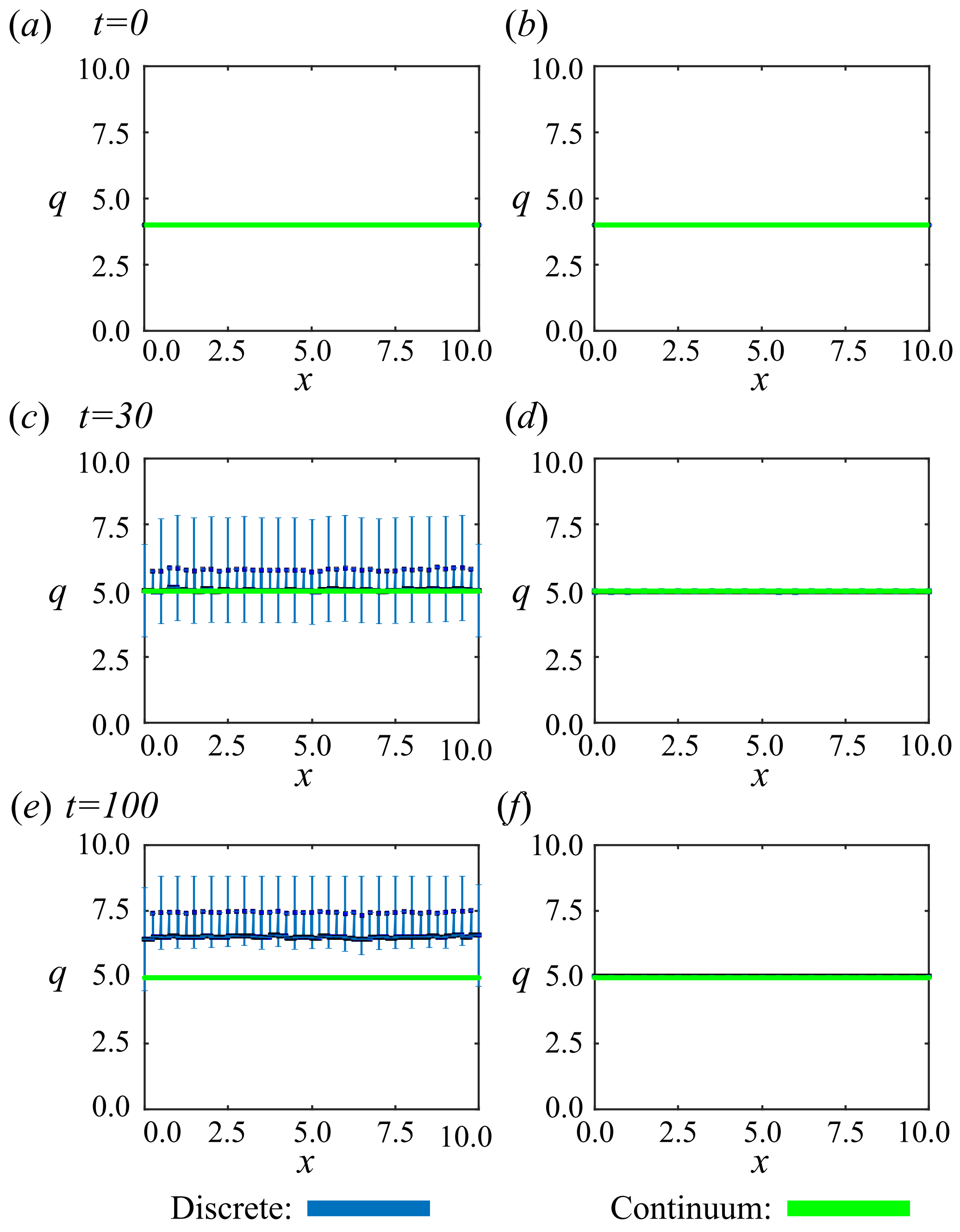


Figure S6: Homogeneous population with piecewise proliferation mechanism. With slow mechanical relaxation, $k = 0.0001$, and faster mechanical relaxation, $k = 1000$, shown in left and right columns, respectively. (a)-(f) Density snapshots at times $t = 0, 30, 100$ where the average and standard deviation (blue error bars) of 2000 discrete simulations are compared to solution of continuum model (green).

In Figure 7 in the main manuscript we show that at later times the solution of the continuum model can differ significantly from the solution to the discrete model with slow mechanical relaxation, $k = 0.0001$, and matches extremely well when $k = 1000$. We now show $N(400)$ for intermediate values of $k$ in Figure S7.

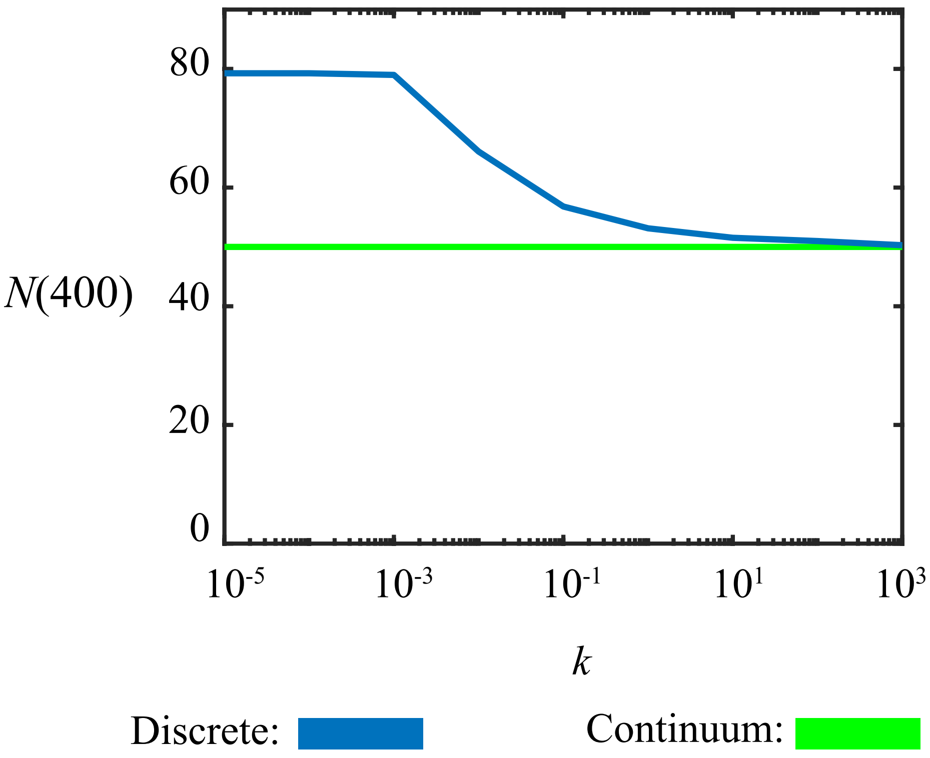


Figure S7: Homogeneous population with piecewise proliferation mechanism. $N(400)$ with varying mechanical relaxation rates through cell stiffness $k$. Increasing the cell stiffness $k$ improves matching between discrete and continuum model. Cell stiffness plotted on a logarithmic axis.

### S4 Mechanical cell competition

We now present additional results for a heterogeneous population: an exact calculation for cell size at mechanical equilibrium, and results for logistic proliferation and death.

#### S4.1 Two populations: cell size at mechanical equilibrium

Here we show that, for cells in extension, cells with lower cell stiffness are larger than cells with higher cell stiffness at mechanical equilibrium. As mechanical relaxation is chosen to be fast relative to the proliferation and death rates the system will be at mechanical equilibrium, except for the short transition following a proliferation or death event.

In Section 3.2 of the main manuscript, we are interested in two adjacent populations which we denote tissue 1 and tissue 2. We assume there are $N_i$ cells in tissue $i$ and cells in tissue $i$ have cell stiffness $K_i$ and resting cell length $A_i$. From previous work (Murphy et al. 2019) the interface position at mechanical equilibrium, $\mathcal{S} = \lim_{t\to\infty} s(t)$, is

$$\mathcal{S} = \frac{\frac{K_1 A_1}{K_2} + \frac{L}{N_2} - A_2}{\frac{K_1}{K_2 N_1} + \frac{1}{N_2}}. \tag{45}$$

Assuming $A_i = 0$ this simplifies to

$$\mathcal{S} = \frac{L}{\frac{K_1 N_2}{K_2 N_1} + 1}. \tag{46}$$

Letting $l_i$ be the length of a cell in tissue $i$ then

$$\begin{aligned} l_1 &= \frac{S}{N_1}, \\ l_2 &= \frac{L - S}{N_2}. \end{aligned} \tag{47}$$

Substituting Equation (46) into Equation (47) gives

$$\begin{aligned} l_1 &= \frac{L}{\frac{K_1 N_2}{K_2} + N_1}, \\ l_2 &= \frac{L}{N_2 + \frac{K_2 N_1}{K_1}}. \end{aligned} \tag{48}$$

It can then be shown that if $K_1 < K_2$ then $l_1 > l_2$. This corresponds to cells of lower stiffness being larger at mechanical equilibrium and this is independent of the number of cells in tissue 1 and 2.

#### S4.2 Mechanical cell competition: logistic combination

In this section, we repeat the scenario presented in Section 3.2 but now with the logistic proliferation and death mechanisms rather than the linear proliferation and death mechanisms. We observe similar qualitative results.

With mechanical relaxation and proliferation only, we observe that the interface position is on average the same as the initial condition which was chosen as mechanical equilibrium (Figure S8). This is expected as proliferation is independent of cell length and therefore mechanical relaxation. Therefore we also expect that the total number of cells in tissue 1 and tissue 2 are, on average, equal (Figure S8i). From Equation (46) we expect the initial condition of mechanical equilibrium to be maintained. Similarly, in the continuum model both tissues have the same proliferation rate therefore the interface position, $s(t)$, is maintained (Figure S8j).

For mechanical relaxation with proliferation and death, we now observe that the cancer cells extend and the healthy cells are smaller. The smaller healthy cells then eventually die and the cancer cells take over the domain. This is because the death mechanism is now dependent on cell length and therefore dependent on mechanical relaxation (Figure S4.2). Note that the total cell number for each population does not decrease below zero, the error bars represent the standard deviation about the mean.

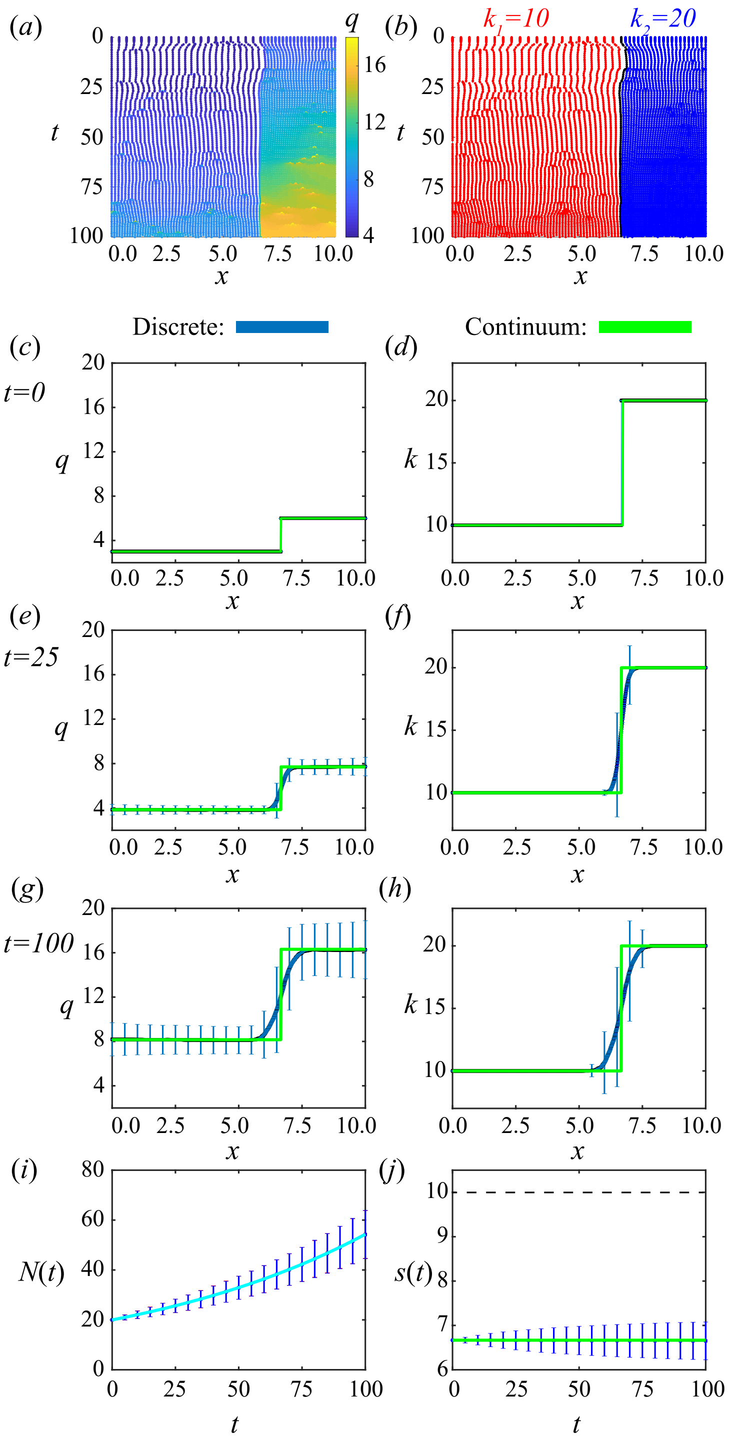


Figure S8: Results for cancer invasion with adjacent populations using **logistic** proliferation and death mechanisms with **proliferation only**. (a),(b) A single realization of cell boundary characteristics for $0 \leq t \leq 100$. Colouring in (a),(b) represent cell density and cell stiffness, respectively. (c)-(d), (e)-(f), (g)-(h) Density and cell stiffness snapshots, left and right, respectively, at times $t = 0, 25, 100$, respectively. (i) Total cell number, $N(t) > 0$, for cancer (red/magenta) and healthy cells (blue/cyan) for the discrete/continuum solutions. (j) Interface position, $s(t)$, where the dotted line shows the edge of the domain. The average and standard deviation (blue error bars) of 2000 discrete simulations are compared to the solution of the continuum model (green).

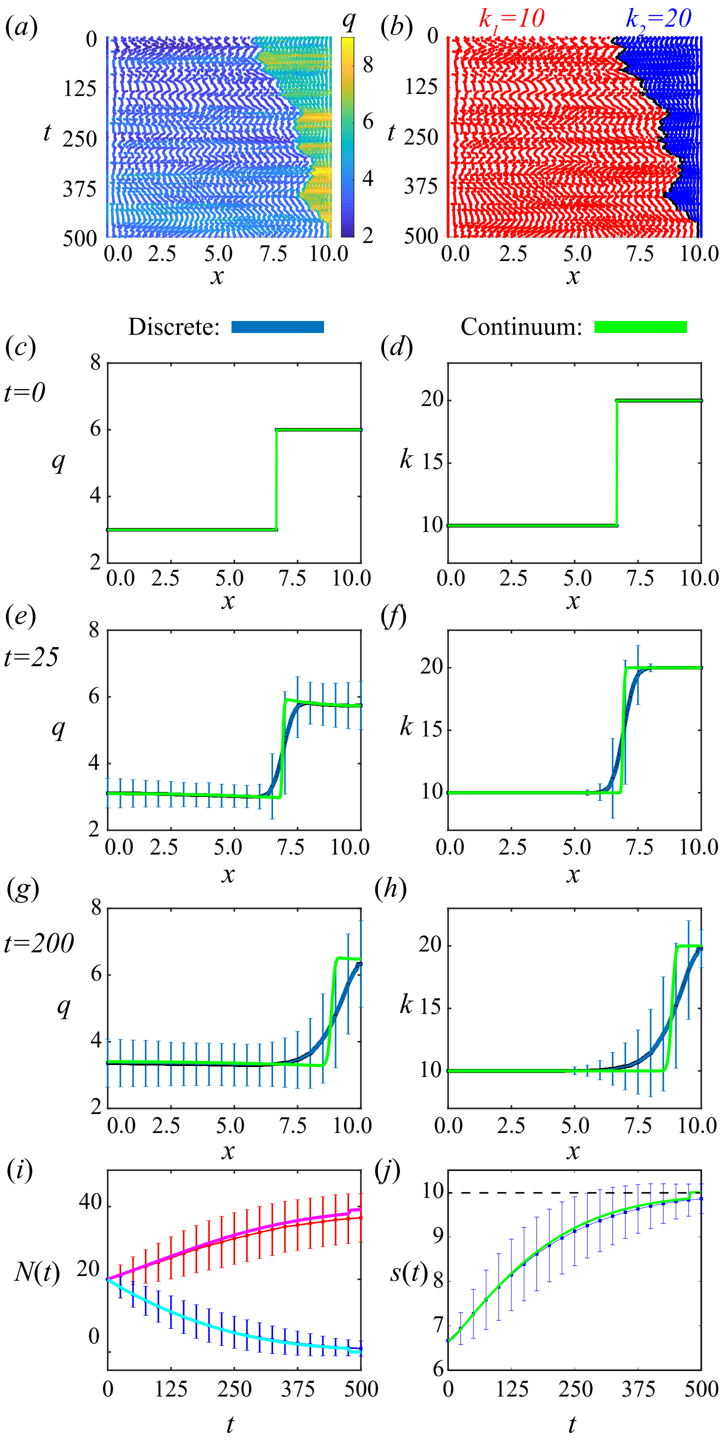


Figure S9: Results for cancer invasion with adjacent populations using **logistic** proliferation and death mechanisms with **proliferation and death**. First row shows a single realization of cell boundary characteristics for $0 \leq t \leq 500$. Colouring in (a),(b) represent cell density and cell stiffness, respectively. (c)-(d), (e)-(f), (g)-(h) Density and cell stiffness snapshots, left and right respectively, at times $t = 0, 25, 200$, respectively. (i) Total cell number, $N(t) > 0$, for cancer (red/magenta) and healthy cells (blue/cyan) for the discrete/continuum solutions. (j) Interface position, $s(t)$, where the dotted line shows the edge of the domain. The average and standard deviation (blue error bars) of 2000 discrete simulations are compared to the solution of the continuum model (green).

##### S4.3 Varying mechanical relaxation rate

In Figure 6 in the main manuscript, mechanical cell competition is considered with cell stiffnesses $K_1 = 10, K_2 = 20$, and linear proliferation and death mechanisms. Here, in Figure S10, we present the same problem with reduced mechanical relaxation rates and observe that the continuum model is a reasonably good approximation even for cell stifnesses $K_1 = 0.0001, K_2 = 0.0002$.

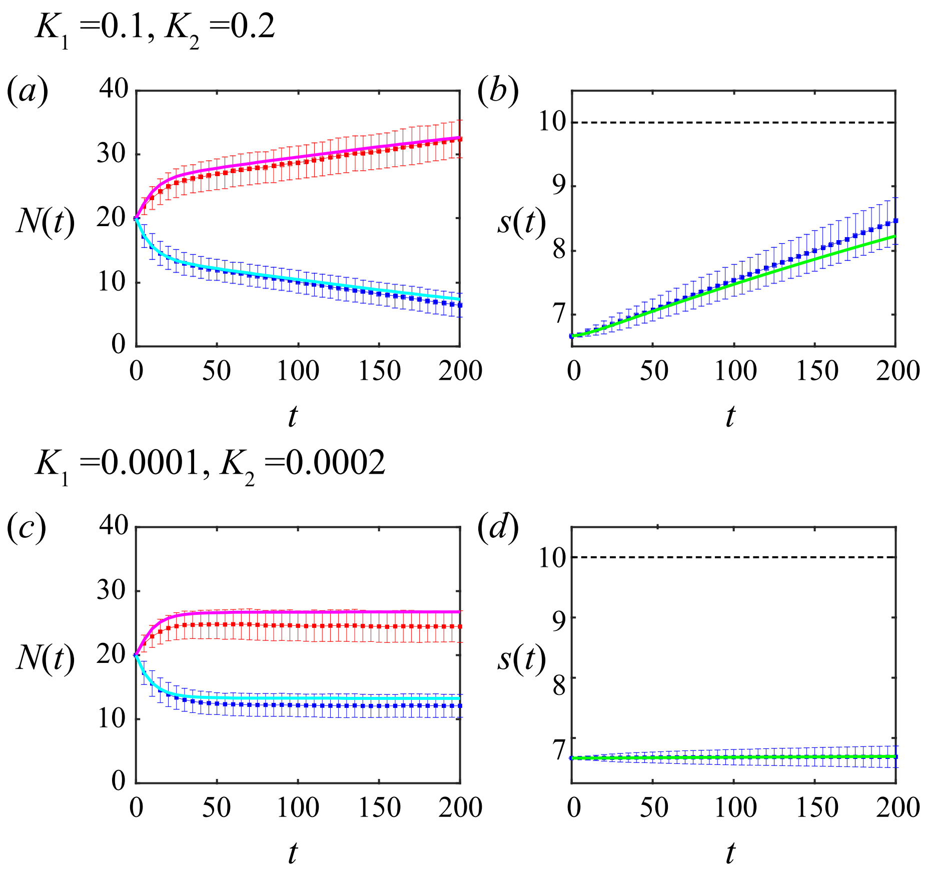


Figure S10: Results for cancer invasion with adjacent populations using **linear** proliferation and death mechanisms with **proliferation and death** with slower mechanical relaxation, $k = 0.1$ in (a,b) and $k = 0.0001$ in (c,d). (a,c) Total cell number, $N(t) > 0$, for cancer (red/magenta) and healthy cells (blue/cyan) for the discrete/continuum solutions. (b,d) Interface position, $s(t)$, where the dotted line shows the edge of the domain. The average and standard deviation (blue error bar) of 2000 discrete simulations are compared to the solution of the continuum model (green).
